# Skeletal muscle nuclei in mice are not post-mitotic

**DOI:** 10.1101/2022.10.24.513426

**Authors:** Agnieszka K Borowik, Arik Davidyan, Frederick F Peelor, Evelina Voloviceva, Stephen Doidge, Matthew P Bubak, Christopher B Mobley, John J McCarthy, Esther E Dupont-Versteegden, Benjamin F Miller

**Author notes:** Corresponding Author: Benjamin F Miller, Aging and Metabolism Research Program, Oklahoma Medical Research Foundation, 825 NE 13th Street, Oklahoma City, Oklahoma 73104.

## Abstract

The skeletal muscle research field generally accepts that nuclei in skeletal muscle fibers (i.e., myonuclei) are post-mitotic and unable to proliferate. Because our deuterium oxide (D_2_O) labeling studies showed DNA synthesis in skeletal muscle tissue, we hypothesized that resident myonuclei can replicate *in vivo*. To test this hypothesis, we used a mouse model that temporally labeled myonuclei with GFP followed by D_2_O labeling during normal cage activity, functional overload, and with satellite cell ablation. During normal cage activity, we observed deuterium enrichment into myonuclear DNA in 7 out of 7 plantaris (PLA), 6 out of 6 tibialis anterior (TA), 5 out of 7 gastrocnemius (GAST) and 7 out of 7 quadriceps (QUAD). The average fractional synthesis rates (FSR) of DNA in myonuclei were: 0.0202 ± 0.0093 in PLA, 0.0239 ± 0.0040 in TA, 0.0076 ± 0. 0058 in GAST, and 0.0138 ± 0.0039 in QUAD, while there was no replication in myonuclei from EDL. These FSR values were largely reproduced in the overload and satellite cell ablation conditions although there were higher synthesis rates in the overloaded PLA muscle. We further provided evidence that myonuclear replication is through endoreplication that results in polyploidy. These novel findings contradict the dogma that skeletal muscle nuclei are post-mitotic and open potential avenues to harness the intrinsic replicative ability of myonuclei for muscle maintenance and growth.

**Graphical Abstract:** 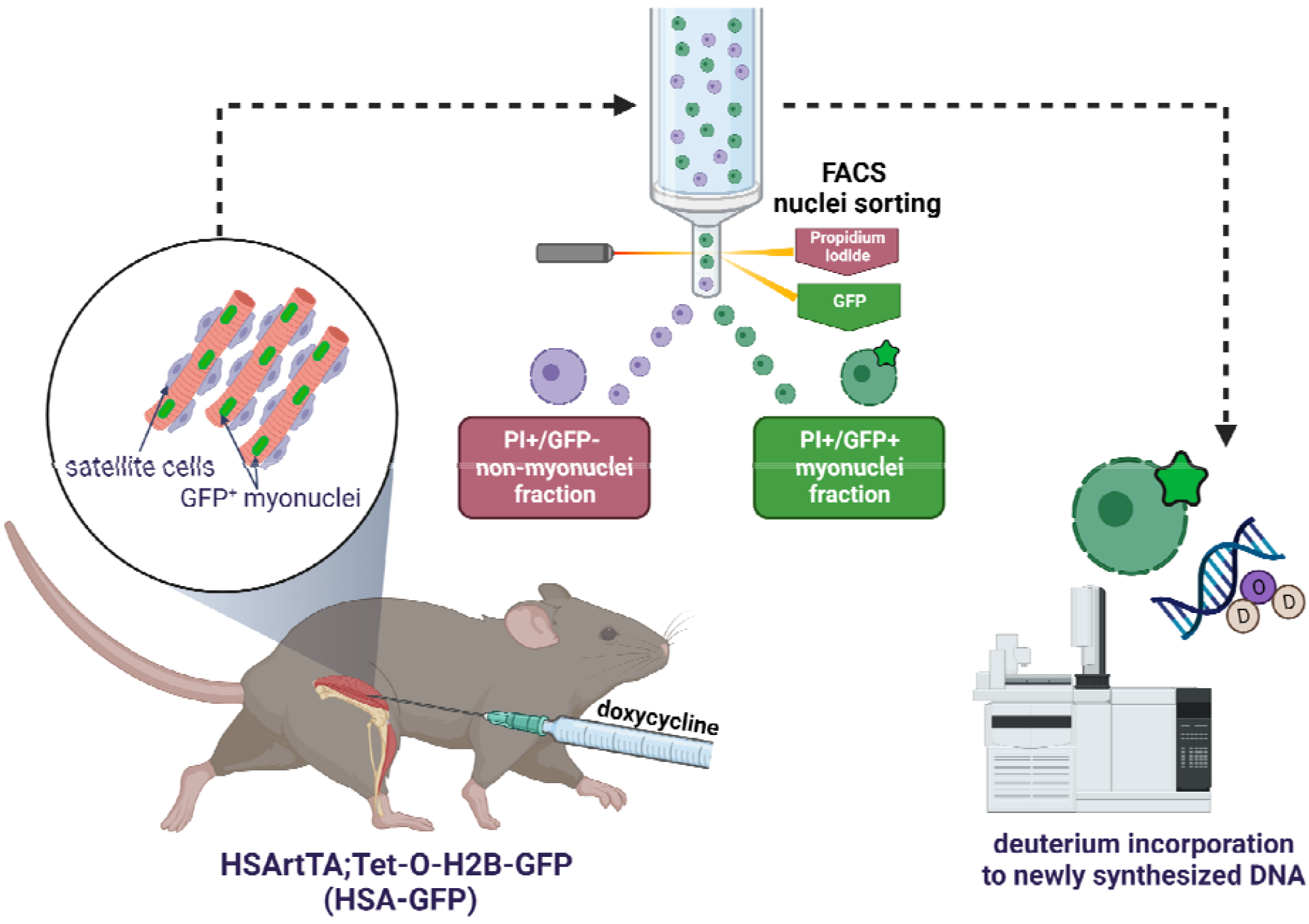

## Introduction

Skeletal muscle is the largest organ in the body by mass and has important locomotive, metabolic, and secretory functions. In addition, skeletal muscle serves as a protein reservoir that can be used for energy metabolism and the maintenance of protein content of tissues during stressful conditions associated with disease and aging^1^. Therefore, preserving muscle mass and function is critically important to maintain health, quality, and duration of life.

The nucleotypic theory states that DNA content is a primary determinant of cell size^2^. The volume of cytoplasm serviced by DNA is referred to as the karyoplasmic ratio, or when applied to skeletal muscle, the resident myonuclear domain^3^. Skeletal muscle myofibers (myocytes) require multiple nuclei (myonuclei) to support a cytoplasmic volume that is five orders of magnitude greater than most cell types^4^. The general consensus in the skeletal muscle research field is that upon terminal differentiation myonuclei enter into a permanent post-mitotic state and are no longer able to replicate their DNA^5^. This perspective became entrenched with the discovery that MyoD was capable of inducing cell cycle arrest independent of differentiation^6^. Therefore, by this paradigm, myocytes are post-mitotic, and the addition of nuclear material requires fusion of satellite cells.

Despite the wealth of data supporting the concept that myonuclei are post-mitotic, there are instances in which myonuclei have been reported to undergo replication. For example, in the newt, skeletal muscle involves the de-differentiation of myocytes into myoblasts which then re-enter the cell-cycle and replicate. Further, terminally differentiated C2C12 myotubes can de-differentiate and re-enter the cell cycle when induced by extracellular factors like inducible SV40 large T antigen oncogene^7^, cyclin D1 and CDK4^8^. In mice, the ablation of satellite cells does not compromise myonuclei abundance or function over time^9–15^, suggesting that muscles have satellite cell-independent capacity to maintain nuclei number. Additionally, myonuclear content does not change and muscular adaptation is not diminished during life-long aerobic exercise of satellite cell-depleted mice^13,16–20^. Studies that have induced high rates of satellite cell myofiber fusion over time did not have the expected increase of myonuclear number, suggesting the existence of myonuclear turnover^12,21^. Finally, although controversial, cardiomyocytes, which are another muscle cell type thought to be post-mitotic, have now been shown to replicate DNA independent of progenitor cells^22^.

The lack of definitive evidence that myonuclei can replicate may stem from the methodological challenges involved in capturing rare replication events. When a rapidly dividing mononuclear cell undergoes nuclear replication, karyokinesis is followed by cytokinesis, which can normally be captured by microscopic techniques. However, the spatial arrangement of contractile elements in a multi-nucleated myofiber does not allow for cytokinesis events. Second, traditional DNA labeling methods like BrdU incorporation indicate that DNA has replicated, but it does not distinguish between a satellite cell or myonucleus as the source of new DNA within a myofiber. Further, because BrdU incorporation requires immunohistochemistry for detection, there can be ambiguity when signals are weak, or determining the meaning of low versus high signals. Finally, myonuclear replication events are likely infrequent, making it challenging to capture a replication event with traditional snapshot approaches.

Deuterium oxide (D_2_O) labeling is a validated approach for measuring DNA synthesis in multiple cell and tissue types^23,24^. D_2_O labeling has distinct advantages over BrdU labeling in that it is incorporated into DNA through *de novo* pathways, not salvage pathways^24^, which minimizes or completely eliminates labeling from DNA repair processes. Further, at the enrichment levels typically used, D_2_O has no known toxicities, does not affect proliferation rates, and is non-mutagenic^24^. Finally, it is simple to administer deuterium as D_2_O in the drinking water of animals for weeks (or months), which is an important advantage when capturing a metabolic event, like myonuclear replication, that may occur at a low rate.

Through our studies that have used D_2_O labeling, we have consistently demonstrated that skeletal muscle has measurable rates of DNA synthesis *in vivo*^25,26^, that these rates are reduced during growth restriction^27,28^, and the rates are increased with mechanical stimulation of muscle^29^. In fact, all studies in which we have isolated DNA from skeletal muscle have found detectable rates of DNA synthesis. Although intriguing, our previous measurements were made on muscle homogenates that included several cell types, including replicative mononuclear cell types, which could have contributed to the measured rates of DNA synthesis. In the current study we first tested whether satellite cells were the source of DNA replication. We then sought to test *in vivo* if the source of DNA replication is skeletal muscle resident myonuclei. To do so, we used a genetic approach that allowed temporal fluorescence labeling of myonuclei followed by D_2_O labeling of DNA replication. We applied these approaches during regular cage activity, with mechanical overload stimulus induced by synergist ablation (SA), and in the absence of satellite cells that may “unmask” myonuclear replicative potential. We hypothesized that skeletal muscle resident myonuclei can replicate DNA *in vivo*. Further, we hypothesized that DNA replication would not be accompanied by mitosis (i.e. endoreplication,) because of the spatial constraints and syncytial nature of myofibers, and therefore would result in polyploidy. Our findings challenge the dogma that skeletal muscle resident myonuclei are post-mitotic and thus fundamentally changes the understanding of the basic biology of skeletal muscle.

## Materials and Methods

### Generation of mouse models

All animal procedures were conducted in accordance with institutional guidelines for the care and use of laboratory animals and were approved by the Institutional Animal Care and Use Committees at the Oklahoma Medical Research Foundation (OMRF) and University of Kentucky. Mice were housed in the vivarium at the OMRF or University of Kentucky under the care of the Department of Comparative Medicine. Animals were group housed (maximum 5 per cage) with *ad libitum* access to food and water in a room on a 14:10 h light:dark cycle with constant temperature and humidity control. Mouse models used for breeding to generate the models, breeding strategies, and genotyping are provided in **Supplementary Methods and Supplementary Tables 1-3**.

### Model validation

Mouse models were validated prior to experimentation. The Pax7-DTA mouse was previously validated by our group and used to deplete adult skeletal muscle of satellite cells ^9,10,15^. For HSA-GFP mice, we performed immunohistochemistry (IHC) analyses on tibialis anterior (TA) muscle cryosections. TAs were isolated from induced mice, embedded in optimal cutting temperature compound (OCT compound), and snap frozen in isopentane^30^. Ten-micron thick longitudinal sections were air dried for 1 hour at room temperature and fixed with 4% PFA for 7 min. Sections were washed with 1x phosphate buffered saline (PBS), and endogenous peroxidases were blocked for 7 minutes with 3% hydrogen peroxide in PBS. We performed epitope retrieval in sodium citrate (10 mM, pH 6.5) for 20 min at 92°C, followed by TrueBlack Autofluorescence Quencher (Biotium) supplemented with Mouse-on-Mouse (MoM) IgG blocking reagent (Vector Laboratories, Burlingame, CA, USA). Sections were washed in PBS and incubated for 15 minutes in streptavidin and biotin (Vector Laboratories, Burlingame, CA, USA). Afterwards, slides were incubated overnight with primary antibodies diluted in 1% TSA blocking reagent, followed 24 hour later with a PBS rinse and secondary antibodies (**see Supplementary Table 4**). The slides were then incubated for 20 min in AlexaFlour555 Tyramide Reagent (1:200, Thermo Fisher Scientific), stained with DAPI (1:10,000 in PBS, Thermo Scientific) for 10 min, and mounted with VectaShield fluorescent mounting media. The images were captured at 20× magnification using both brightfield and polarized light on a confocal microscope (Zeiss LSM-710, Oberkochen, Germany) with subsequent analysis using ImageJ (U. S. National Institutes of Health, Bethesda, Maryland, USA). During described analysis assessors were blinded to the experimental groups.

### Study procedures

#### Satellite cell ablation experiment

When we started the current studies, we proposed that satellite cells could be the source of our measured DNA synthesis rates in skeletal muscle tissue and that age may change these synthesis rates. We hypothesized that satellite cell replication decreased with age and that ablating satellite cells reduced or eliminated any measured DNA synthesis rates. To test these hypotheses, 5 and 12-month-old female Pax7-DTA mice were given i.p. injections of either vehicle (15% ethanol in sunflower seed oil, +SC) or tamoxifen (2 mg/day in sunflower seed oil, -SC) for five consecutive days (**Figure 1A**). At 6 months (Young) or 24 months (Old) mice were randomized to either sedentary control or 6 weeks of voluntary exercise, to stimulate satellite cells. Exercised mice were singly housed, and we monitored running distance daily (ClockLab software, Actimetrics, Wilmette, IL). Sedentary mice were housed in identical cages without wheels. To measure DNA synthesis rates, we gave the mice an i.p. injection of sterile 99% deuterium oxide (D_2_O) and 4% D_2_O-enriched drinking water for the 6-week sedentary or exercise period according to our previously published methods^31,32^.

**Figure 1:**
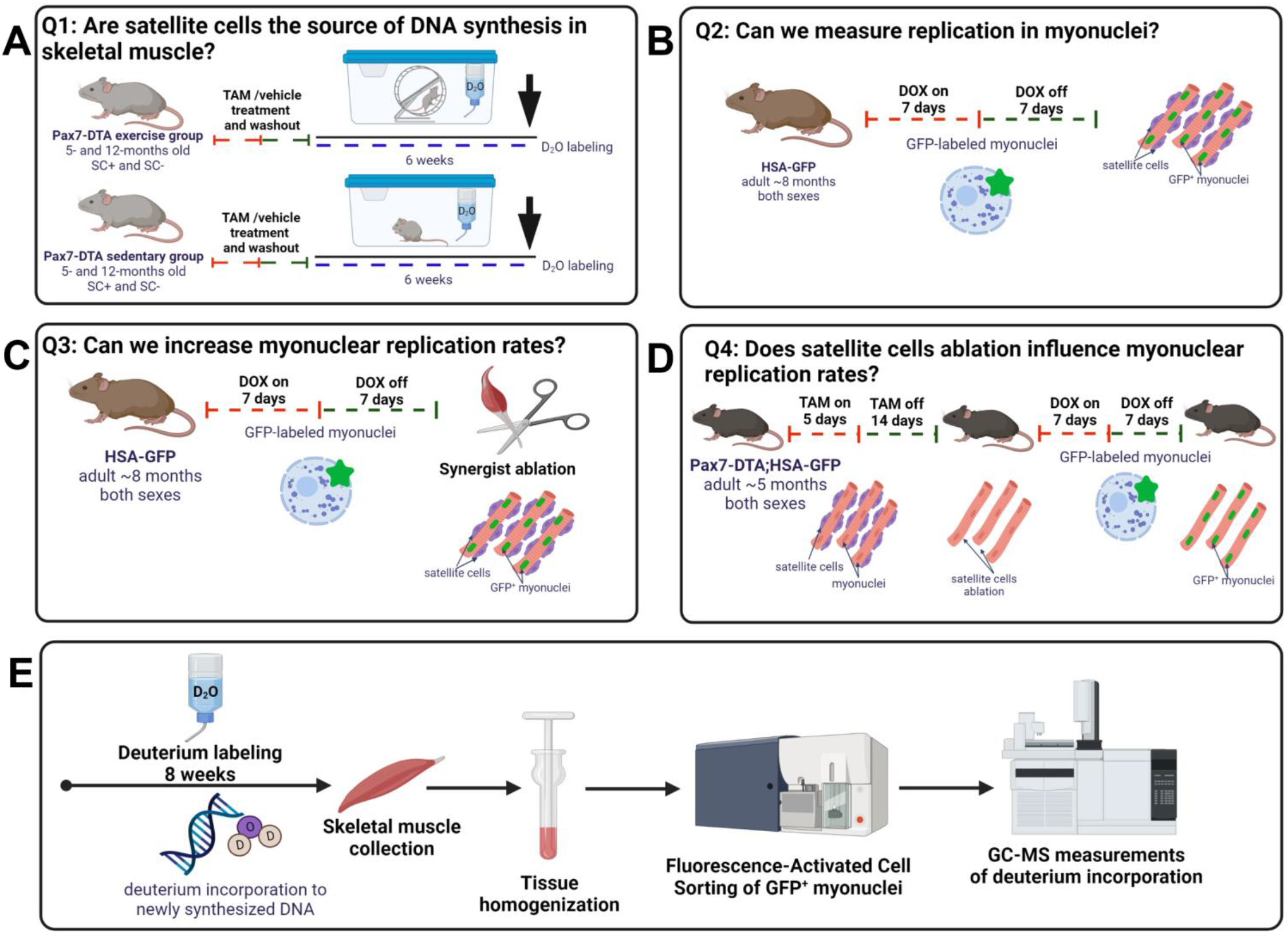
Overview of the experimental design. **(A)** To identify the source of measured DNA synthesis in skeletal muscle, we used Pax7-DTA mice. Animals underwent tamoxifen (TAM) or vehicle treatment to induce conditional ablation of satellite cells. After washout, mice were assigned into sedentary or exercise group (wheel running) for 6 weeks. Mice were labeled with the stable isotope deuterium oxide (D_2_O) for the entire time of the intervention. **(B)** To test if skeletal muscle resident myonuclei possess the ability to replicate *in vivo*, we used the HSA-GFP mouse model, which uses doxycycline (DOX) treatment for temporal labeling (Tet-ON) of resident myonuclei with GFP. **(C)** To test if myonuclear replication can be stimulated by mechanical overload, DOX-treated HSA-GFP mice were subjected to synergist ablation (SA) surgery. **(D)** To test if satellite cells (SCs) depletion stimulates myonuclei to replicate, we used the Pax7-DTA;HSA-GFP mouse model, which uses TAM treatment to deplete SCs and DOX treatment to label resident myonuclei with GFP. In myonuclei replication experiments after removing DOX, and a washout period, we provided D_2_O for 6-10 weeks to measure DNA synthesis in sorted GFP^+^ (resident myonuclei) and GFP^-^ (non-myonuclei) fractions by GC-QQQ.

#### Myonuclei replication

We then performed a series of experiments to determine if resident myonuclei were the source of the replicating DNA in skeletal muscle tissue (**Figure 1B-E**). Our strategy “pre-labeled” myonuclei with GFP, and then determined if GFP^+^ myonuclei (i.e., resident myonuclei) accumulated deuterium-labeled DNA. To accomplish this approach, we induced GFP-labeling in resident myonuclei of HSA-GFP mice using 0.5 mg/ml doxycycline (DOX) in drinking water with 2% sucrose (for taste) (**Figure 1B**). After 7 days we stopped DOX to turn off H2B-GFP labeling, therefore “pre-labeling” myonuclei. We then performed a 7-day washout period to ensure all DOX was cleared from the mice with no additional myonuclear H2B-GFP labeling, including any cells that could potentially be in a transition state. Following this washout, we provided an i.p. injection of 99% D_2_O followed by 8% D_2_O-enriched drinking water to label newly synthesized DNA. D_2_O labeling continued for 6-10 weeks in the normal activity group, and 8-10 weeks in the mechanical overload and satellite cell-ablation groups (**Figure 1C&D**), after which we harvested the following skeletal muscles: plantaris (PLA), extensor digitorum longus (EDL), tibialis anterior (TA), gastrocnemius (GAST) and quadriceps (QUAD) muscles. From these individual muscles we isolated GFP^+^ skeletal muscle resident myonuclei using fluorescence-activated cell sorting (FACS). Last, we analyzed the sorted GFP^+^ myonuclei for D_2_O incorporation into DNA using GC-QQQ (**Figure 1E**). We used this strategy for the following three experiments with any deviations noted.

To determine if resident myonuclear replication occurs normally without an intervention, we labeled seven-month-old HSA-GFP mice (n=7) during normal cage activity (**Figure 1B**). To test if mechanical loading increased rates of resident myonuclei replication, we performed synergist ablation surgery (SA)^10,33^ to overload the PLA muscle in seven-month-old HSA-GFP mice (n=5). The SA was performed immediately after DOX treatment (**Figure 1C**) according to previous procedures^10,33^. In brief, to remove the GAST and soleus, we performed a longitudinal incision on the dorsal aspect of the lower hind limb under isoflurane anesthesia. The tendon of the GAST muscle was isolated and guided the excision of the GAST and soleus muscle without disturbing the nerve, blood supply and PLA muscle. The incisions were sutured, and the next day mice received their i.p. bolus injection to start the D_2_O labeling period. Our last experiment sought to determine if ablation of satellite cells stimulated resident myonuclei replication because of the loss of a stem cell population. We first used five days of i.p. tamoxifen treatment (20 mg/mL dissolved in corn oil) in five-month-old Pax7-DTA;HSA-GFP mice (n=7) to ablate satellite cells (**Figure 1D**). After a 7-day washout, the mice followed the same GFP and D_2_O labeling protocol as the other experiments.

### Tissue collection

Mice were euthanized with CO_2_ inhalation followed by cervical dislocation. Blood was collected and then centrifuged at 2000 *g* for 10 min at 4°C. Plasma was then aliquoted and frozen at −80°C for further analyses. Muscles from both hindlimbs were immediately harvested, trimmed of fat and connective tissues, and further processed for nuclei isolation. Bone marrow was isolated from the femur and tibia to determine the deuterium enrichment of a fully turned over DNA pool. In some mice, we collected the liver for a positive control for FACS sorting to determine potential polyploidy.

### Nuclear isolation

Nuclear isolation was performed on fresh tissue according to the modified procedure previously described by von Walden *et al*.^34^. We pooled hindlimb muscles of the same type from the left and right limb to increase yield of nuclear number. Muscles were placed in 1 mL homogenization buffer containing 5 mM PIPES, 85 mM KCl, 1 mM CaCl_2_, and 5% sucrose supplemented with 2x protease inhibitors (Halt™ Protease Inhibitor Cocktail, Thermo Scientific) and 0.25% NP-40 prepared in ddH_2_O (pH =7.2). Tissue was minced into small pieces using microscissors, then transferred to the manual Dounce homogenizer (glass on glass), where nuclei were liberated using 15-20 strokes. The muscle suspension was diluted in 15 mL of buffer, then passed through 70 μm filter on top of the 50 mL conical tube. The crude nuclear suspension was further diluted with an additional 5 mL of buffer, passed through 40 μm filter, then centrifuged for 6 minutes at 3,000 RPM at 4°C with a swinging-bucket rotor. The supernatant was aspirated, the crude nuclear pellet was re-suspended in 2 mL of buffer without NP-40 and transferred to a FACS tube. The crude nuclear suspension was spiked with 6 μL of propidium iodide (PI, Sigma, 1mg/mL) to label all nuclei, then subjected to FACS analysis.

### Fluorescence activated cell sorting FACS

Propidium iodide positive (PI^+^)/GFP^+^ nuclei (myonuclei) and PI^+^/GFP^-^ nuclei (non-myonuclei) fractions were sorted using a Moflo XDP (Beckman-Coulter, Indianapolis, IN, USA) equipped with a Sapphire 488 laser, Coherent Cube 640 laser, Coherent Cube 405 laser and analyzed using FlowJo software (FlowJo, LLC, Ashland, OR, USA). Total nuclei population was defined based on PI signal. The myonuclei and non-myonuclear fractions were classified after elimination of doublets via forward scatter area versus height. To prevent nuclear membrane rupture, nuclei were sorted with 60 PSI pressure, using a 70 μm nozzle. Sorted fractions were collected in 1x phosphate buffered saline (PBS) supplemented with 1x protease inhibitors (Halt™ Protease Inhibitor Cocktail, Thermo Scientific) and stored at -80°C until further analysis. For the autofluorescence and background normalization during sorting, we used the control sample from the untreated HSA-GFP mouse, where myonuclei are not GFP-labeled. Nuclear polyploidy was measured as a fluorescence intensity of PI (area under the curve). Nuclei isolated from hepatocytes were used as an internal genome size control.

### DNA isolation

In our initial Pax7-DTA experiment, total DNA was extracted from approximately 15 mg of muscle tissue and 300 mg of bone marrow using the QiAmp DNA mini kit (Qiagen, Valencia, CA). In subsequent experiments, nuclear DNA was isolated from the sorted myonuclear and non-myonuclear fractions using QIAmp DNA Micro Kit (Qiagen, Germantown, MD, USA) with adjusted reagents’ volumes to the samples size. DNA from bone marrow was isolated using QiAamp DNA mini kit (Qiagen, Germantown, MD, USA). Resultant DNA concentrations were determined using a NanoDrop (Thermo Fisher Scientific).

### Body water enrichment

To determine the deuterium body water enrichment, samples were prepared and analyzed according to our previous published methods for GC-MS analysis (initial satellite cell ablation experiment)^35^ or with a liquid water isotope analyzer (Los Gatos Research, Los Gatos, CA, USA)^32,36^.

### Analysis of deuterium enrichment into DNA

Isolated DNA from muscle tissue, bone marrow, or cells were analyzed for deuterium enrichment as we have previously described^32,36–38^. Briefly, isolated DNA was hydrolyzed overnight at 37°C with nuclease S1 and potato acid phosphatase. Hydrolysates were reacted with pentafluorobenzyl hydroxylamine and acetic acid and then acetylated with acetic anhydride and 1-methylimidazole. Dichloromethane extracts were dried, resuspended in ethyl acetate, and analyzed by GC–QQQ (Agilent 8890 GC coupled to Agilent 7010B GC-QQQ, Santa Clara, CA, USA) on a DB-17 column with negative chemical ionization, using helium as carrier and methane as the reagent gas. The fractional molar isotope abundances at m/z 435 (M0, mass isotopomer) and 436 (M1) of the pentafluorobenzyl triacetyl derivative of purine deoxyribose were quantified using ChemStation software. Excess fractional M+1 enrichment (EM1) was calculated as follows:

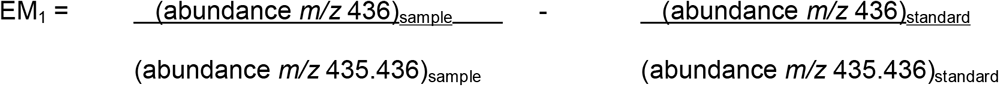

where sample and standard refer to the analyzed sample and unenriched pentafluorobenzyl triacetyl purine deoxyribose derivative standard, respectively. To control for background enrichments, we used two procedures. In the first, we used unenriched pentafluorobenzyl triacetyl purine deoxyribose derivative standard at increasing concentrations to control for mass shifts with increasing DNA concentration. In the second, we used tissue from unlabeled mice prepared side-by-side with our labeled mice. We also performed two calculations of synthesis using bone marrow from the same animal, which represents a fully turned over population of cells, or plasma D_2_O enrichment with mass isotopomer distribution analysis (MIDA) adjustment^31,32^. These additional analyses reduced the chances of having a false positive of deuterium enrichment. The fraction new was divided by time (in days) to obtain a fractional synthesis rate (FSR) in %/day.

### Myosin Heavy Chain (MyHC) determination (Fiber typing)

The muscles from both hindlimbs were quickly removed, trimmed of excess fat and connective tissues and flash-frozen in liquid nitrogen.

Muscles were cut at the mid-belly and the distal half was mounted for tissue sectioning in a cryostat to determine muscle fiber cross-sectional area and for immunohistochemical analyses. Seven-μm thick sections were used in immunolabeling experiments to demonstrate fiber type by probing for the major myosin heavy chain (MHC) isoforms in mice skeletal muscle as previously described^39^. Briefly, frozen sections were incubated overnight with isotype specific anti-mouse primary antibodies for myosin heavy chain (MyHC) I (1:75, IgG2B, BA.D5), MyHC IIa (supernatant, IgG1, SC.71), and MyHC IIx (supernatant, IgM, 6H1) from DSHB, along with laminin (1:100, Sigma-Aldrich, St. Louis, MO, USA). Sections were then incubated with secondary antibodies (1:250, goat anti-mouse IgG2B alexa fluor 647; 1:500, IgG1 alexa fluor 488; 1:250, IgM alexa fluor 555) from Invitrogen, all diluted in PBS, along with the secondary antibody for laminin (1:150, IgG AMCA, Vector). Sections were mounted with VectaShield mounting medium (Vector), and images were captured at 20x. Fiber type distribution was quantified using Myovision as percent MyHC I, IIa, IIx, and IIb (no staining)^40^. For all immunohistochemistry analyses we counted a minimum of 600 fibers for each animal and assessors were blinded to the experimental groups.

### Statistical approach

For the initial Pax7-DTA experiment we used a two-factor ANOVA (vehicle/tamoxifen × sedentary/exercised) to determine if satellite cell ablation impacted DNA replication rates in muscle tissue. If a significant interaction was detected, Scheffé post-hoc comparisons were performed to identify the source of significance with p ≤0.05. The remaining myonuclear DNA replication experiments required a different approach. Because we were testing if a previously undescribed phenomenon could occur, our statistical analyses were designed to test the yes/no conclusion of each hypothesis. For each sample, we determined zero enrichment by two approaches. If enrichment was greater than zero for that run, we consider the sample to have measurable DNA synthesis. *A priori* we expected that for some trials myonuclear DNA replication may not occur. A non-observation in a single trial did not mean that DNA replication cannot occur. To eliminate a potential spurious finding, we required that a single positive observation of DNA replication was confirmed in at least one other mouse. If two or more mice were positive, we considered the hypothesis supported. In our actual experiments, we confirmed DNA replication early, and performed additional studies to establish the rate at which this replication was occurring.

## Results

Our initial hypothesis was that satellite cells were the source of DNA synthesis in skeletal muscle tissue. Therefore, we used the Pax7-DTA mouse model to determine if satellite cell ablation eliminated DNA replication. We performed these experiments in young and older mice, and during sedentary or exercise conditions. Although there was a significant increase in DNA replication with exercise, satellite cell ablation did not decrease DNA synthesis rates (**Figure 2**). Since the ablation of satellite cells did not account for DNA synthesis rates that we consistently measure in skeletal muscle *in vivo*, we sought an approach to directly measure resident myonuclei replication.

**Figure 2:**
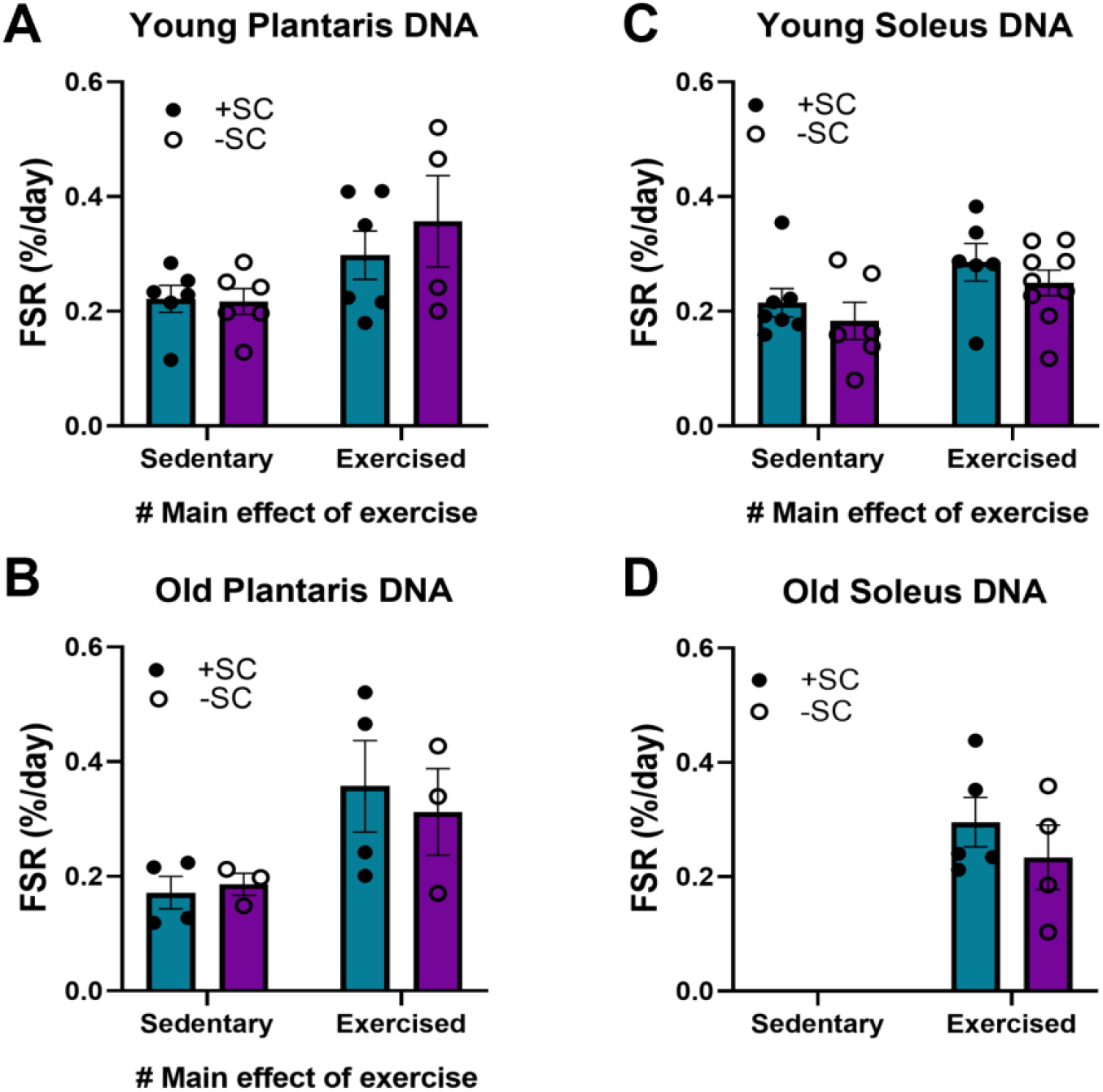
Impact of satellite cells (SC) ablation on Fractional Synthesis Rate (FSR) of DNA in plantaris (A,B) and soleus (C,D). FSR (%/day) was determined in plantaris and soleus muscles of young and old control and exercised mice. A two-way ANOVA (satellite cell presence by exercise) determined a significant effect of exercise, which increased FSR of DNA. However, there were no main effects of satellite cell ablation. Teal boxes = control (+SC); purple boxes = ablated (-SC). Values expressed as mean ± SEM.

To test that resident myonuclei can replicate DNA, we used the HSA-GFP mouse, which uses DOX treatment for temporal and specific fluorescent tagging (Tet-ON) of residential myonuclei with GFP^30^. Importantly, after turning off the labeling, we included a 7-day washout period prior to starting D_2_O labeling to make sure our D_2_O labeling did not catch a satellite cell in a transition state of fusing to the myofiber and donating a nucleus that may have just undergone DNA replication. We repeated the validation of the HSA-GFP mouse model for effective GFP labeling of myonuclei as previously performed by our group^30^. We visualized fibers in longitudinal section (**Figure 3A**) and single fibers (**Figure 3B**). Our staining showed that Pax7+ nuclei (pink) never co-localized with GFP-positive myonuclei (green) within laminin border (purple). After we confirmed the GFP labeling by IHC, we proceeded to FACS analysis on isolated nuclei. We aimed to be restrictive in our gating to minimize the chance of false positives. Before each FACS analysis, we set the gates and autofluorescence signals with control samples from the non-induced HSA-GFP mice spiked with propidium iodide (PI). In the first step, we used forward and side scatter parameters to visualize the density of crude nuclear fraction from the muscle tissues (**Figure 4A**). Afterward, we set up a PI-signal gate to positively select the population of total nuclei from the tested sample (**Figure 4B**). The nuclei within the PI-positive gate were further analyzed for the presence of GFP fluorescence (**Figure 4C**). The same gating strategy was applied for sorting samples coming from the induced HSA-GFP mice (**Figure 4D-F**). In our approach, we distinguished myonuclei as PI^+^/GFP^+^ population, while nuclei coming from other cell types (non-myonuclei) were PI^+^/GFP^-^. To further confirm the purity of sorted myonuclear fraction, we visualized nuclei isolated from muscle tissue before and after FACS analysis using fluorescence microscopy (**Figure 4G**). A representative quantification of the sorting efficiency is presented in **Supplementary Figure 1**. Both myonuclear and non-myonuclear fractions were saved for further analysis of deuterium incorporation into DNA.

**Figure 3:**
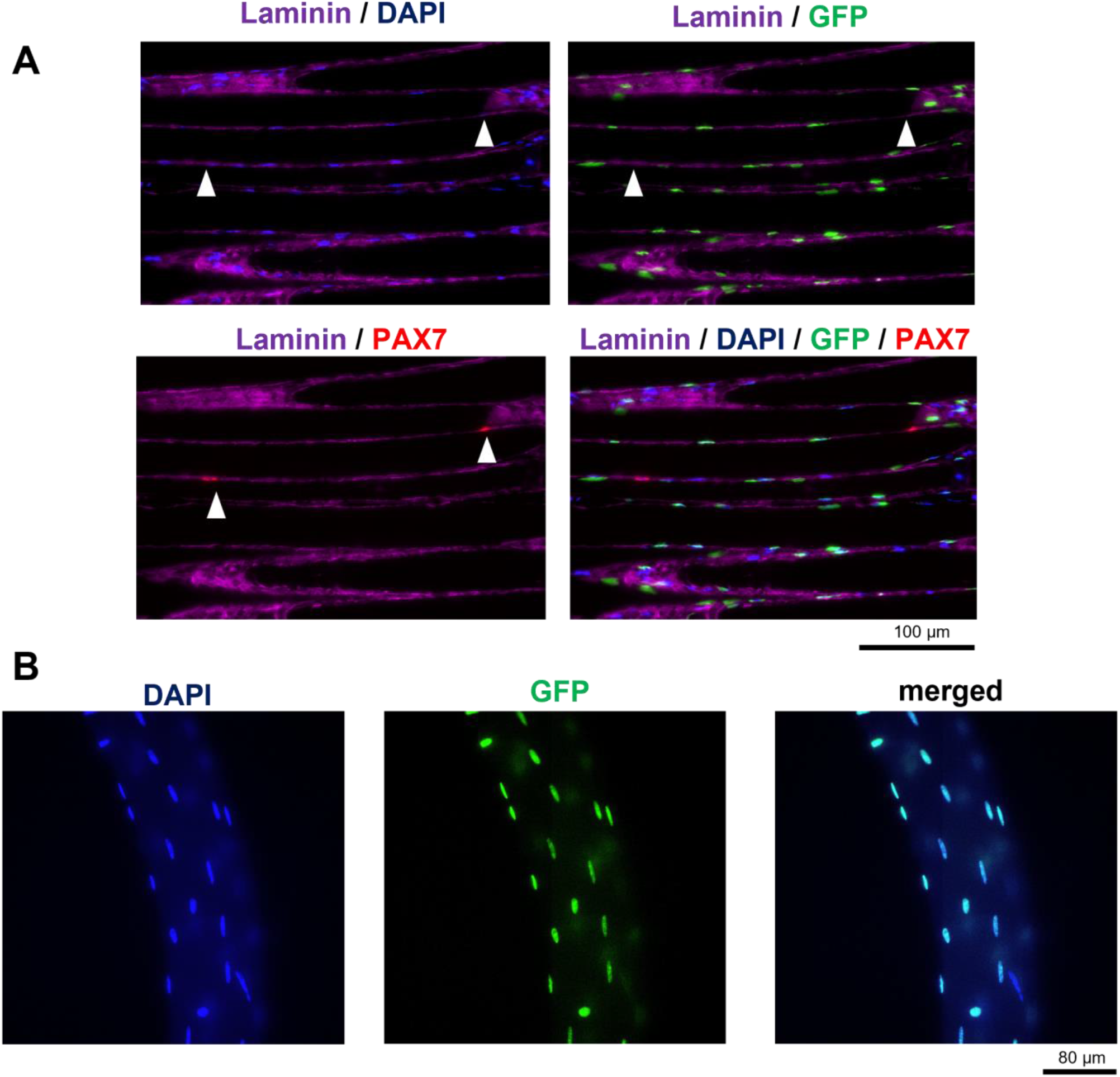
Representative microscopic images of the validation of the HSA-GFP skeletal muscle after DOX treatment. (**A)** Longitudinal sections of TA muscle from HSA-GFP mouse. DAPI = total nuclei, GFP = myonuclei, PAX7 = satellite cells. GFP signal from myonuclei doesn’t overlap with PAX7 signal from satellite cells. Scale bar = 100μm. (**B)** Single fiber images of EDL muscle from HSA-GFP mouse with GFP-labeled myonuclei. Scale bar = 80μm.

**Figure 4:**
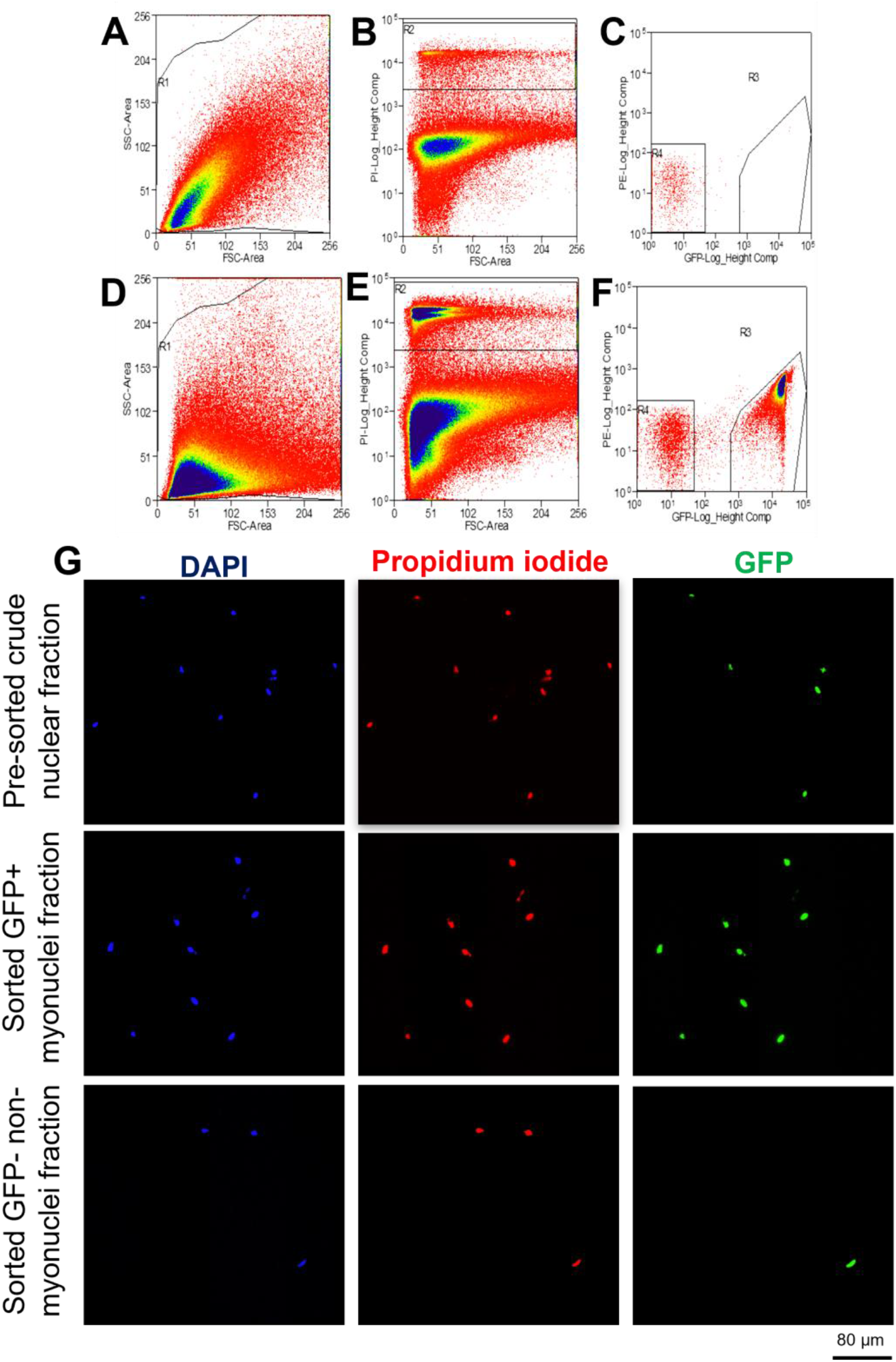
Fluorescence activated cell sorting (FACS) gating strategy for myonuclei and non-myonuclei sorting from skeletal muscle homogenates. Upper panel = gating strategy for muscle homogenates from non-induced HSA-GFP control mice. Lower panel = gating strategy for muscle homogenates from DOX-treaded HSA-GFP mice. **(A**,**D)** Scatterplot of forward scatter (area) by side scatter (area) of muscle homogenate. **(B**,**E)** Scatterplot of propidium iodide (PI, height) by forward scatter (area). Gate R2 indicates total nuclear population. **(C**,**F)** Scatterplot of propidium iodide (PI, height) by GFP (height). Gate R3 indicates myonuclei (PI^+^/GFP^+^); gate R4 indicates non-myonuclei (PI^+^/GFP^-^). **(G)** Representative images of pre-sorted crude nuclear fraction; sorted myonuclear fraction (PI^+^/GFP^+^) and sorted non-myonuclear fraction (PI^+^/GFP^-^). Scale bar = 80μm.

We analyzed the nuclear yield of GFP^-^ and GFP^+^ fractions to compare to previously published values. Myonuclei made up 55-65% of total sorted nuclei from the muscle homogenates from normal cage activity (**Figure 5A**), overload (**Figure 5B**), and satellite cell ablation (**Figure 5C**) groups. There was one exception in the overloaded PLA where only 35% of the nuclei were myonuclei (**Figure 5B**), likely due to the infiltration of inflammatory cell types. We next assessed DNA synthesis rates (%/day) of the GFP^-^ and GFP^+^ fractions of several muscles from multiple animals for each condition. We performed our calculation using two methods to establish a zero enrichment to not have a false positive. Here we report the values from the bone marrow correction, whereas the body water with MIDA correction is presented in **Supplementary Tables 5-16**. In each condition, the first couple of mice confirmed evidence of myonuclear replication. Therefore, we added more mice in each group to establish how consistent this phenomenon was. During normal cage activity, we observed deuterium enrichment into myonuclear DNA in 7 out of 7 PLA, 6 out of 6 TA, 5 out of 7 GAST and 7 out of 7 QUAD. The average FSR of replicating myonuclei was: 0.0202 ± 0.0093 in PLA, 0.0239 ± 0.0040 in TA, 0.0076 ± 0. 0058 in GAST, and 0.0138 ± 0.0039 in QUAD, while EDL appeared not to have replicative myonuclei although the GFP^-^ fraction did (**Figure 5D**). SA increased PLA mass by approximately 53% (**Supplemental Figure 2**). The FSR of myonuclei in the mechanical overloaded PLA was 0.1361 ± 0.0311 (**Figure 5E**) while the other muscles did not seem to change from normal cage activity (0.0328 ± 0.0103 in TA, 0.0145 ± 0. 0045 in QUAD). Again, EDL muscles appeared to not replicate DNA. Finally, we examined myonuclear replication in muscle in which satellite cells were ablated. We expected that stem cells depletion would unmask myonuclear replicative potential and stimulate it. Surprisingly, the FSR of resident myonuclei appeared not to change with satellite cell ablation (0.0019 ± 0.0030 in EDL, 0.0129 ± 0.0057 in PLA, 0.0172 ± 0.0059 in TA, 0.0107 ± 0. 0040 in GAST, and 0.0125 ± 0.0075 in QUAD) (**Figure 5F**). When qualitatively comparing resident myonuclei to other cell types (i.e. the GFP^-^ fraction), the GFP^-^ fraction had higher rates of synthesis across all conditions. The synthesis values were remarkably consistent across conditions and muscles except for the overloaded PLA (**Figure 5D, E, and F**). Finally, to calculate the percentage of the total nuclei (GFP^-^ and GFP^+^) that had new DNA within the labeling period, we multiplied the FSR for each muscle by the number of sorted nuclei from that tissue, and then divided it by total sorted nuclei. We found that around 0.7-1% of myonuclei replicated during the labeling period and that number increased to ∼3% in overloaded PLA muscles (**Figure 5G, H and I**).

**Figure 5:**
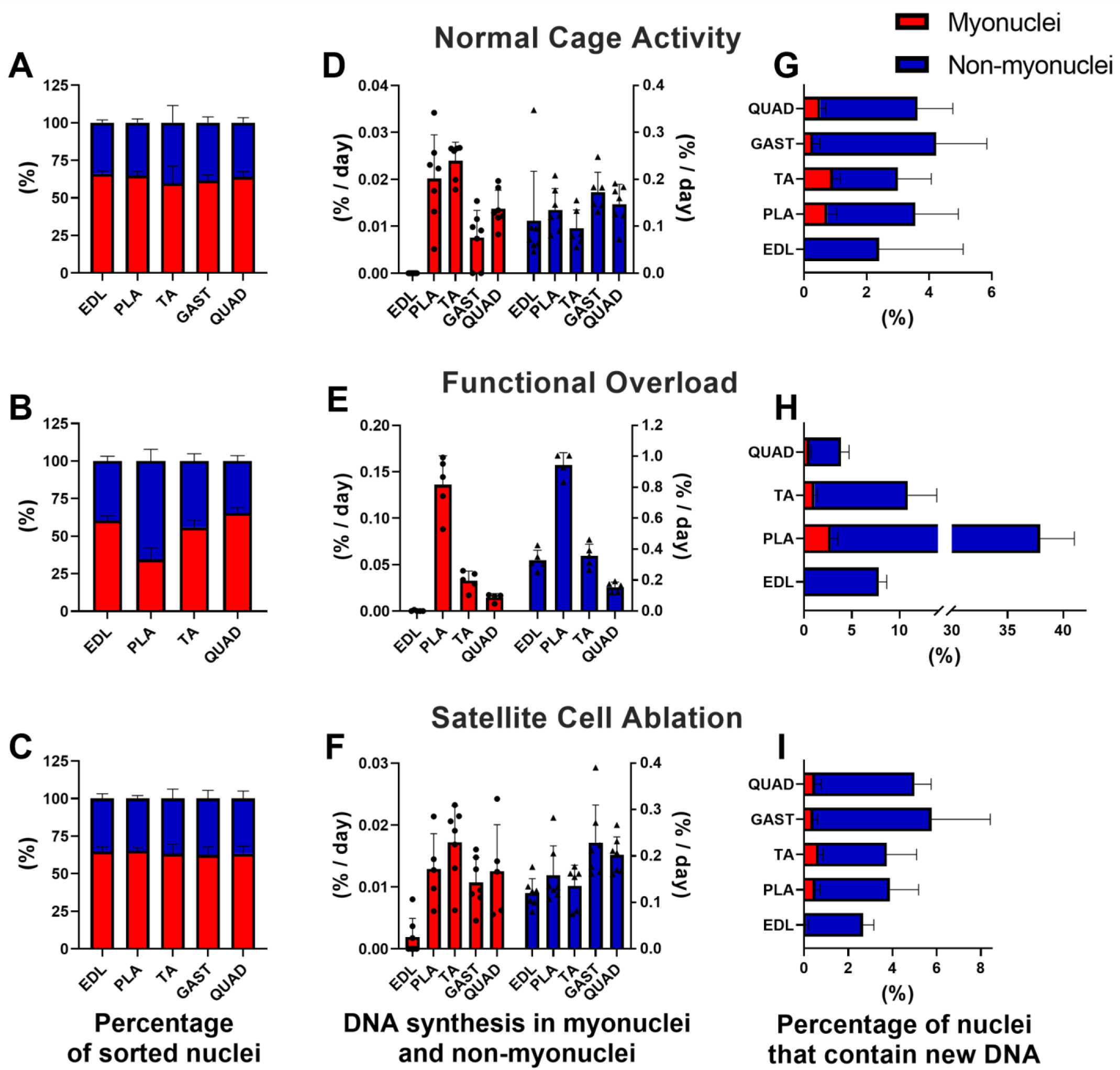
DNA proliferation rates in sorted myonuclear and non-myonuclear fractions. **(A-C)** Nuclei sorting yields from skeletal muscles homogenates. **(D-F)** DNA fractional synthesis rates (FSR, %/day) in myonuclei and non-myonuclei fractions. **(G-I)** Percentage of nuclei that contain new DNA within labeling time in myonuclei and non-myonuclei fractions. Red boxes = myonuclei fraction; blue boxes = non-myonuclei fraction.

It was intriguing to us that EDL had no replication in GFP^+^ nuclei, so we fiber typed our muscles to determine if fiber type differences could explain the results. From our analyses (**Supplemental Figure 3**) there was nothing strikingly different about fiber-type composition between muscles that could explain this finding.

After acquiring evidence of DNA synthesis in resident myonuclei, we wanted to determine if the DNA synthesis resulted in polyploidy given that this is a well-described phenomenon in other tissues like liver and heart^41,42^. Therefore, we isolated nuclei from livers to use as an internal genome size control (**Figure 6A**). We show representative results for the GFP^+^ myonuclei isolated from PLA muscle during normal cage conditions (**Figure 6B**), after functional overload (**Figure 6C**), and after satellite cells ablation (**Figure 6D**, other muscles are shown in **Supplemental Figure 4-6**). Small peaks identifying genome duplications are visible within diploid to hexaploid range. However, myonuclei from the overloaded PLA had more distinct peaks indicative of tetraploidy. The relatively small peaks align with the observation that 1-3% of myonuclei were replicatively active.

**Figure 6:**
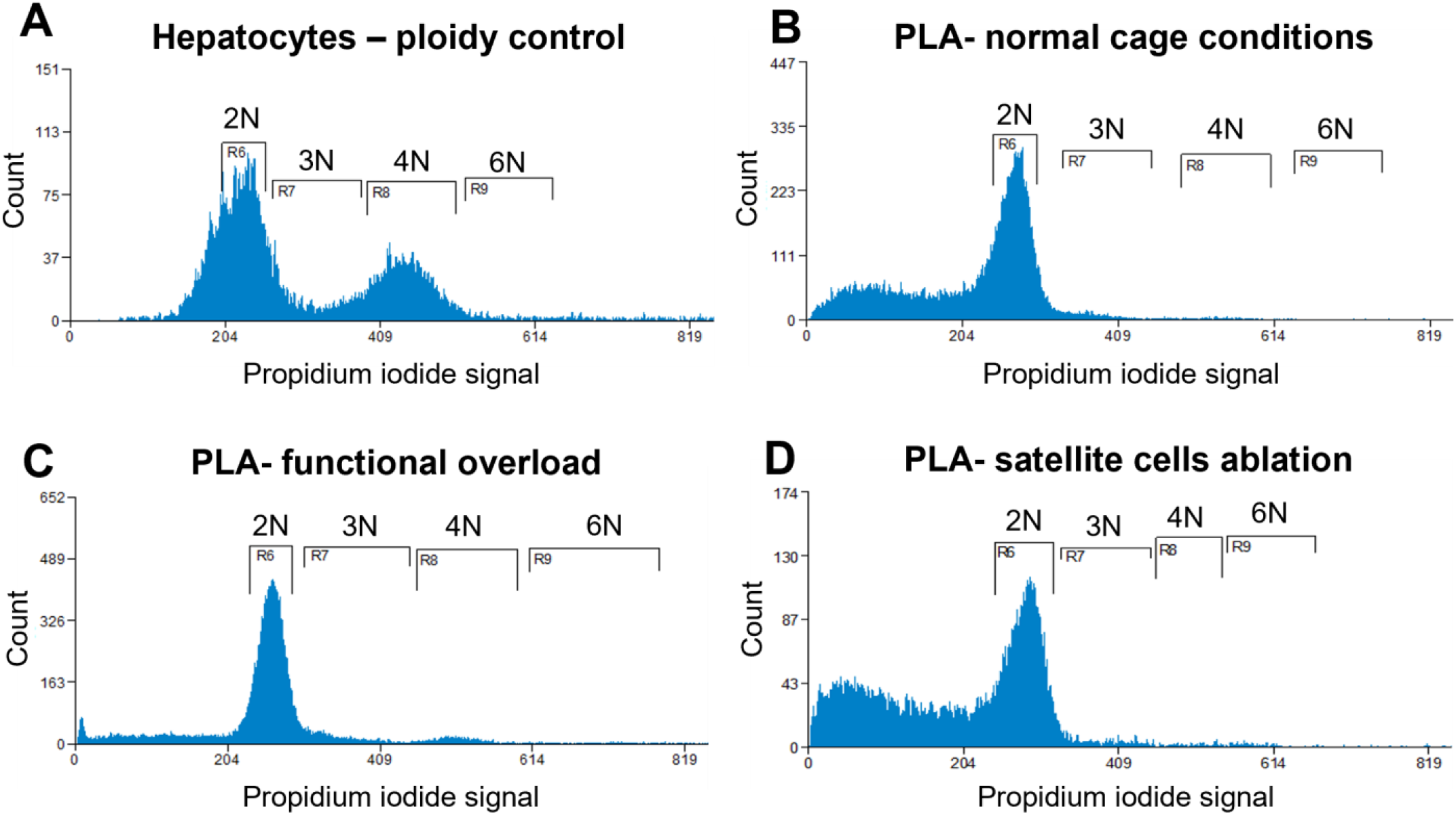
Representative analysis of ploidy levels in sorted myonuclei from plantaris muscles. Cell flow cytometry of hepatocytes **(A)**, myonuclei isolated from plantaris skeletal muscles from animals kept in the normal cage conditions **(B)**, mice after functional overload **(C)**, mice after satellite cells ablation **(D)**. The peaks corresponding to diploid nuclei are labeled 2N, triploid nuclei as 3N, tetraploid nuclei as 4N and hexaploid nuclei as 6N.

## Discussion

We sought to test if DNA in resident myonuclei can replicate. We first ruled out that satellite cells were the source of DNA synthesis we found in previous studies of skeletal muscle tissue. We then used temporal genetic labeling of myonuclei followed by D_2_O labeling to demonstrate that there is DNA synthesis in myonuclei that varies by muscle. We also showed that the rates of resident myonuclei DNA synthesis are higher during overload, but not in the absence of satellite cells. Finally, we demonstrated that DNA synthesis was consistent with endoreplication and resultant polyploidy.

Because these findings challenge current dogma regarding the post-mitotic nature of myonuclei, we took several steps to demonstrate the rigor of our approach. These findings create a new area of investigation where resident myonuclei could be a therapeutic target for preventing muscle loss, increasing muscle growth, and improving clinical conditions.

*DNA synthesis in muscle is not from Pax7*^*+*^ *cells*. In 2011, we first demonstrated that there was DNA synthesis in human skeletal muscle^25^. We also found that aerobic exercise increased the rate of DNA synthesis and that this increase in DNA synthesis correlated with increases in muscle protein synthesis^25^. We speculated that satellite cell replication was the cause of the DNA synthesis including the higher levels that occurred with exercise. Since that study, we have reported changes in DNA synthesis in skeletal muscle with such models as growth-restricted mice^27,28^, and with mechanical stimulation of muscle^29^. Again, we reasoned that changes in satellite cell replication could account for the changes in DNA synthesis. Therefore, in the current study, we ablated satellite cells with the rationale that DNA synthesis would decrease or go to zero in the absence of satellite cells. Contrary to our initial hypothesis, there were no changes in DNA synthesis with satellite cell ablation in multiple muscles of young and old animals at rest or with exercise training. This finding led us to propose that nuclei in other cell types were replicating DNA. We considered potential approaches of systematically ablating one cell type at a time to directly measure a specific cell type. Since we were intrigued by previous indirect evidence of myonuclei replication, we designed a strategy to unambiguously test if myonuclei can replicate DNA.

### Myonuclear DNA synthesis and muscle specificity

The most important finding of this study was that under all conditions studied there was DNA synthesis in mammalian myonuclei. This finding is the first direct measurement of DNA replication in mammalian resident myonuclei *in vivo*, thus establishing the precedent that adult myofibers are not completely post-mitotic as previously thought. These results build upon previous findings from other lower organisms such as the newt where myofibers de-differentiate into myoblasts and then proliferate^43^. They also confirm findings from studies *in vitro*, that myocytes can be dedifferentiated and provoked to re-enter the cell cycle^44,45^. Finally, they show that myonuclei replication is not just a property of cardiomyocytes^42,46,47^ but also skeletal myocytes.

We were able to compare DNA synthesis rates in resident myonuclei to a mix of nuclei from other cell types within the muscle because we isolated GFP^+^ and GFP^-^ fractions. The rates of DNA synthesis in resident myonuclei were about 10x less than the pooled sample of other cell types. Therefore, in our previous studies on muscle tissue, only about 10% of the DNA synthesis can be attributed to myonuclei as opposed to other mononuclear cell types such as pericytes, endothelial cells, fibroblasts, satellite cells and immune cells. There is muscle specificity in the DNA synthesis rates: the EDL had essentially no DNA synthesis under any of the conditions examined, whereas the TA had relatively high rates of DNA synthesis compared to the other muscles. It is tempting to speculate that fiber-type might influence the ability of a myonucleus to replicate DNA; however, our fiber typing revealed no major difference in the fiber-type composition of the EDL or TA that might account for the observed differences in the rate of DNA synthesis. Unfortunately, we did not have a muscle type that was predominantly Type I fibers since a pooled soleus sample had insufficient nuclear number for our analyses. We therefore speculate that the degree of muscle activation determines the rates of replication.

SA surgery had a clear effect on DNA synthesis in the overloaded PLA muscle. SA is a well-established mechanical overload model that can double mass over 14 days^10^, whereas in the current study it increased approximately 53% over 8 weeks. It is likely that the 8-week-period had an initial rapid growth period followed by a normalization of growth. SA surgery is known to be accompanied by damage, inflammation, and swelling in the initial days^33,48^. Because of this damage, there is potentially some muscle regeneration beyond just muscle growth, thus complicating the interpretation. Nonetheless, it is interesting that the muscle overloaded by the SA surgery, the PLA, had a large increase in DNA synthesis whereas the other muscles did not. In addition, the other lower limb muscles (EDL and TA) approximately doubled DNA synthesis in the non-myonuclear fraction, whereas the myonuclear fraction of the EDL and TA did not. It is likely that the non-myonuclear fraction was capturing inflammatory cell types. These changes during overload also lead to the intriguing question of whether the resident myonuclei replication facilitated the rapid growth of the PLA. More studies with a physiological overload stimulus are needed to explore this question.

Interestingly, satellite cell ablation appeared to have no impact on DNA synthesis rates in the myonuclear fraction of multiple muscles. We hypothesized that removing satellite cells would take away a likely contributor to maintaining the myonuclei pool and would thus increase myonuclear replication to maintain myonuclear number. A further confirmation of our model is that the non-myonuclear fraction did decrease DNA synthesis rates with satellite cell ablation. This finding leads us to believe that under conditions of normal cage activity (i.e. non-intervention or non-mechanically stressed) there is little contribution of satellite cells to maintaining the myonuclear pool. These findings confirm previous studies^11^. Future studies should add an additional stimulus to mice with satellite cell ablation to further differentiate the contributions of myonuclear replication and satellite cells to maintaining myonuclear number.

We need to provide one further caution on the measured rates of DNA synthesis. When we initially designed the study, we decided on an 6-10-week D_2_O-labeling period because we were concerned that we were trying to capture rare events. The longer period of labeling would facilitate greater enrichment of the DNA and a better chance for detection on the GC-QQQ. Normal practice of an isotope study is to calculate a rate using the fraction new divided by the period of labeling to get a fractional synthesis rate (%/time). In a recent study, we demonstrated that for some proteins far less than 100% of the proteins turn over^37^. Therefore, at some period, all the proteins that were going to incorporate label do so and the extended period of labeling does not result in a greater fraction new. To illustrate why this is important, suppose a study made its measurement at 8 weeks and determined that 50% of the proteins were new, but those proteins actually reached that 50% by 4 weeks. The 8-week experiment would greatly underestimate the rate of synthesis because 50%/8 weeks = 6.25%/week when the true rate was 50%/4 weeks = 12.5%/week. We provide this example because in unpublished studies of tissues, we have noted that the fraction new of DNA goes up rapidly and then plateaus. This finding leads us to believe that there are pools of DNA that turnover at different rates, with some that do not turn over at all. Therefore, it is possible that the synthesis of DNA underestimates the true rates of myonuclear replication of sub-populations because of the prolonged labeling period. Future studies should incorporate shorter labeling periods to distinguish potential pools of rapid and slowly replicating myonuclei.

### Endoreplication and polyploidy

Endoreplication, a process observed in cardiomyocytes and hepatocytes, serves as a well-known strategy to facilitate cell growth in terminally differentiated cell types in which mitosis would compromise function. Endoreplication can either result in polyploidy, or a rare variant in which mitosis proceeds through nuclear division without cytokinesis leading to multinucleated cells^49^. Endoreplication in skeletal myofibers would facilitate an increase of DNA content without investment of substantial energetic resources to reorganize the contractile elements to facilitate cytokinesis. Endoreplication is also consistent with low to nonexistent tumorigenesis in skeletal muscle since a myocyte would not undergo mitosis. It is possible that endoreplication occurs in resident myonuclei to facilitate the variation in cell size (e.g. hypertrophy) throughout the lifespan. From our data, we have clear indication that the relatively small percentage of myonuclei replicating could be undergoing endoreplication with resultant polyploidy. This finding is particularly apparent in the PLA after SA surgery. It is likely that since the rate of DNA synthesis went up in the PLA with SA surgery, that the amount of polyploidy did too. Although it is known that polyploidy is a strategy to facilitate larger cell size^50^, there is also controversy over whether polyploidy inhibits regenerative potential^51^. Since growth and regeneration are two different programs, with one expanding current cell size while the other invokes a developmental program, it remains to be determined if this polyploidy could be advantageous for normal load-induced muscle growth.

*Is this replication meaningful?* The FSR of non-overloaded muscle is approximately 0.007 - 0.022%/day, which equals 0.049 – 0.154%/week. As discussed, until further studies can be done with shorter labeling periods, this value is likely a minimal rate. Even so, these rates are comparable to 0.08%/week reported for unperturbed cardiomyocytes^52^. Over a year, this means that at minimum 2.5 – 8% of resident myonuclei replicate DNA. As demonstrated, functional overload increased resident myonuclei replication so that FSR was 1%/week. Our sorting procedures were capturing 350,000 to 2 million myonuclei (depending on muscle), which equates to several hundred to thousands of resident myonuclei replicating DNA in each muscle over the period of 6-10 weeks. At present, these are estimations and further studies are needed to understand what may change these rates over time.

However, the most important point is that we demonstrated that myonuclear replication of DNA does occur, and that despite slow rates there are a significant number of myonuclei that replicate DNA. The fact that myonuclear DNA replication occurs *in vivo* leads us to conclude that the process can be driven. If true, the manipulation of myonuclear replication opens several avenues for therapeutic targets independent of stem cell therapies. By this strategy, one could harness the replicative capacity that already exists within the muscle to facilitate muscle maintenance and growth. Similar strategies are already moving forward with cardiomyocytes^53^.

### Rigor of approach

From the outset of the study, we knew that to demonstrate resident myonuclei replication, we would need to be rigorous in our approach by eliminating nuclei from other cell types and the ability to capture potentially rare replication events. First, we validated our mouse model by IHC. Skeletal muscle staining confirmed that DOX treatment induced GFP-labeling of myonuclei, and that satellite cells were not GFP-tagged. Second, we added a washout step after DOX induction to ensure we were not labeling stem cells in transition to myonuclei. Third, we used antibody-independent FACS with a rigorous gating strategy that allowed us to sort myonuclei as a PI^+^/GFP^+^ population, different from the nuclei from the other cell types that were PI^+^/GFP^-^. The success of our gating strategy was demonstrated by fluorescence microscopy images of pre- and post-sorted fractions. Fourth, because we were concerned about the detection of low enrichments on the GC-QQQ, we used two methods, a standard curve and unlabeled myonuclei, to establish background enrichment. With both methods we were able to clearly demonstrate enrichment above background. In addition, we used two precursor enrichments; bone marrow, which represents a fully turned over DNA pool, and body water with MIDA adjustment^24^ to calculate rates. And finally, we used 8-month-old mice to rule out any contribution of developmental programs to myonuclei replication.

When performing studies of DNA synthesis, we are often asked how we know we are not measuring the synthesis of mtDNA or DNA repair processes. In the current study, mtDNA can be ruled out because we specifically isolate nuclei. We can rule out DNA repair processes two ways. Labeling of new DNA is known to come completely from *de novo* deoxynucleoside (dN) synthesis^54^, which is the precursor pool labeled by D_2_O. D_2_O does not label dN salvage pathways, which provides the majority of dN for repair^23^. Further, it is estimated that DNA sustains 10,000-1,000,000 lesions per/cell/day, which equates to approximately <0.0003% of the 2.8 billion base pair (for a mouse) genome/day. Therefore, if all lesions were repaired, there would be a maximum of 0.005% new base pairs/month. Given that repair processes could only label a tiny fraction of DNA, and that DNA repair processes rely on dN salvage pathways^55^, which D_2_O does not label^23^, we would not measure repair processes. Finally, if a previously unrecognized process allowed for deuterium incorporation into DNA through repair processes that relied on de novo-synthesized dN, replicating just one compliment of nuclear DNA (2.8 billion base pairs) would dilute any of the label from repair. Therefore, we can safely rule out that our measured rates of DNA synthesis were in mtDNA or for DNA repair.

## Conclusions

Using a novel strategy relying on temporal genetic labeling and subsequent D_2_O labeling, we unambiguously demonstrated that DNA in myonuclei can replicate. These amounts of DNA synthesis differed by muscle, and changed with mechanical overload, but not satellite cell ablation. It appears that there is endoreplication of resident myonuclei that results in polyploidy.

These findings contradict the dogma that skeletal muscle myocytes are post-mitotic and open potential avenues to harness the intrinsic replicative ability of the cell for muscle maintenance and growth.

## Acknowledgements and Funding

We would like to acknowledge the advice of Dr. Linda Thompson and the staff at the OMRF Cell Sorting Core. We further acknowledge Dr. Kevin Murach (University of Arkansas) for his helpful discussions, Dr. Karyn Hamilton (Colorado State University) for her contributions to the study on the Pax7-DTA mice and initial study design, and Amy Confides for her technical assistances with MyHC analysis. This project was supported by NIH R21AR077387 (BFM, JJM, and EDV) and T32 AG052363 (AD and MPB). Dr. Miller was also supported by the Department of Veteran Affairs (VA I01 BX005592). The content is solely the responsibility of the authors and does not necessarily represent the official views of the National Institutes of Health or the Department of Veterans Affairs. The funding sources played no role in the conduct, writing, or submission of the manuscript for publication.

## References

1. Wolfe RR. The underappreciated role of muscle in health and disease. Am J Clin Nutr. 2006;84(3):475–482.

2. Gregory TR. Coincidence, coevolution, or causation? DNA content, cell size, and the C-value enigma. Biol Rev Camb Philos Soc. 2001;76(1):65–101.

3. Hall ZW, Ralston E. Nuclear domains in muscle cells. Cell. 1989;59(5):771–772.

4. Bruusgaard JC, Liestøl K, Ekmark M, Kollstad K, Gundersen K. Number and spatial distribution of nuclei in the muscle fibres of normal mice studied in vivo. J Physiol. 2003;551(2):467–478.

5. Holtzer H, Marshall JM, Finck H. An analysis of myogenesis by the use of fluorescent antimyosin. J Biophys Biochem Cytol. 1957;3(5):705–724.

6. Weintraub H, Davis R, Tapscott S, et al. The myoD Gene Family: Nodal Point During Specification of the Muscle Cell Lineage. Science (80-). 1991;251(4995):761–766.

7. Endo T, Nadal-Ginard B. Reversal of myogenic terminal differentiation by SV40 large T antigen results in mitosis and apoptosis. J Cell Sci. 1998;111(8):1081–1093.

8. Latella L, Sacco A, Pajalunga D, et al. Reconstitution of Cyclin D1-Associated Kinase Activity Drives Terminally Differentiated Cells into the Cell Cycle. Mol Cell Biol. 2001;21(16):5631–5643.

9. Jackson JR, Mula J, Kirby TJ, et al. Satellite cell depletion does not inhibit adult skeletal muscle regrowth following unloading-induced atrophy. Am J Physiol -Cell Physiol. 2012;303(8).

10. Mccarthy JJ, Mula J, Miyazaki M, et al. Effective fiber hypertrophy in satellite cell-depleted skeletal muscle. Development. 2011;138(17):3657–3666.

11. Fry CS, Lee JD, Mula J, et al. Inducible depletion of satellite cells in adult, sedentary mice impairs muscle regenerative capacity without affecting sarcopenia. Nat Med. 2015;21(1):76–80.

12. Keefe AC, Lawson JA, Flygare SD, et al. Muscle stem cells contribute to myofibres in sedentary adult mice. Nat Commun. 2015;6.

13. Murach KA, Confides AL, Ho A, et al. Depletion of Pax7+ satellite cells does not affect diaphragm adaptations to running in young or aged mice. J Physiol. 2017;595(19):6299–6311.

14. Lee JD, Fry CS, Mula J, et al. Aged Muscle Demonstrates Fiber-Type Adaptations in Response to Mechanical Overload, in the Absence of Myofiber Hypertrophy, Independent of Satellite Cell Abundance. J Gerontol A Biol Sci Med Sci. 2016;71(4):461–467.

15. Englund DA, Murach KA, Dungan CM, et al. Depletion of resident muscle stem cells negatively impacts running volume, physical function, and muscle fiber hypertrophy in response to lifelong physical activity. Am J Physiol Cell Physiol. 2020;318(6):C1178–C1188.

16. Goh Q, Song T, Petrany MJ, et al. Myonuclear accretion is a determinant of exercise-induced remodeling in skeletal muscle. Elife. 2019;8.

17. Jackson JR, Kirby TJ, Fry CS, et al. Reduced voluntary running performance is associated with impaired coordination as a result of muscle satellite cell depletion in adult mice. Skelet Muscle. 2015;5(1):1–17.

18. Fry CS, Lee JD, Jackson JR, et al. Regulation of the muscle fiber microenvironment by activated satellite cells during hypertrophy. FASEB J. 2014;28(4):1654–1665.

19. Englund DA, Figueiredo VC, Dungan CM, et al. Satellite Cell Depletion Disrupts Transcriptional Coordination and Muscle Adaptation to Exercise. Function. 2020;2(1):33.

20. Fry CS, Kirby TJ, Kosmac K, McCarthy JJ, Peterson CA. Myogenic Progenitor Cells Control Extracellular Matrix Production by Fibroblasts during Skeletal Muscle Hypertrophy. Cell Stem Cell. 2017;20(1):56–69.

21. Pawlikowski B, Pulliam C, Betta ND, Kardon G, Olwin BB. Pervasive satellite cell contribution to uninjured adult muscle fibers. Skelet Muscle. 2015;5(1).

22. Bergmann O, Bhardwaj RD, Bernard S, et al. Evidence for cardiomyocyte renewal in humans. Science. 2009;324(5923):98–102.

23. Neese RA, Misell LM, Turner S, et al. Measurement in vivo of proliferation rates of slow turnover cells by 2H2O labeling of the deoxyribose moiety of DNA. Proc Natl Acad Sci U S A. 2002;99(24):15345–15350.

24. Hellerstein MK, Busch R, Neese RA, Awada M, Hayes GM. Measurement of cell proliferation by heavy water labeling. Nat Protoc. 2007;2(12):3045–3057.

25. Robinson MM, Turner SM, Hellerstein MK, Hamilton KL, Miller BF. Long-term synthesis rates of skeletal muscle DNA and protein are higher during aerobic training in older humans than in sedentary young subjects but are not altered by protein supplementation. FASEB J. 2011;25(9):3240–3249.

26. Miller BF, Drake JC, Naylor B, Price JC, Hamilton KL. The measurement of protein synthesis for assessing proteostasis in studies of slowed aging. Ageing Res Rev. 2014;18:106–111.

27. Drake JC, Bruns DR, Peelor FF, et al. Long-lived Snell dwarf mice display increased proteostatic mechanisms that are not dependent on decreased mTORC1 activity. Aging Cell. 2015;14(3):474–482.

28. Drake JC, Peelor FF, Biela LM, et al. Assessment of mitochondrial biogenesis and mTORC1 signaling during chronic rapamycin feeding in male and female mice. J Gerontol A Biol Sci Med Sci. 2013;68(12):1493–1501.

29. Miller BF, Hamilton KL, Majeed ZR, et al. Enhanced skeletal muscle regrowth and remodelling in massaged and contralateral non-massaged hindlimb. J Physiol. 2018;596(1):83.

30. Iwata M, Englund DA, Wen Y, et al. A novel tetracycline-responsive transgenic mouse strain for skeletal muscle-specific gene expression. Skelet Muscle. 2018;8(1):1–8.

31. Drake JC, Bruns DR, Peelor FF, et al. Long-lived crowded-litter mice have an age-dependent increase in protein synthesis to DNA synthesis ratio and mTORC1 substrate phosphorylation. Am J Physiol -Endocrinol Metab. 2014;307(9):E813.

32. Brown JL, Lawrence MM, Borowik A, et al. Tumor burden negatively impacts protein turnover as a proteostatic process in noncancerous liver, heart, and muscle, but not brain. J Appl Physiol. 2021;131(1):72–82.

33. Tsika RW, Hauschka SD, Gao L. M-creatine kinase gene expression in mechanically overloaded skeletal muscle of transgenic mice. Am J Physiol. 1995;269(3 Pt 1).

34. Von Walden F, Rea M, Mobley CB, et al. The myonuclear DNA methylome in response to an acute hypertrophic stimulus. Epigenetics. 2020;15(11):1151–1162.

35. Miller BF, Pharaoh GA, Hamilton KL, et al. Short-term Calorie Restriction and 17α-Estradiol Administration Elicit Divergent Effects on Proteostatic Processes and Protein Content in Metabolically Active Tissues. Journals Gerontol Ser A Biol Sci Med Sci. 2020;75(5):849.

36. Kobak KA, Batushansky A, Borowik AK, et al. An In Vivo Stable Isotope Labeling Method to Investigate Individual Matrix Protein Synthesis, Ribosomal Biogenesis, and Cellular Proliferation in Murine Articular Cartilage. Funct (Oxford, England). 2022;3(2).

37. Abbott CB, Lawrence MM, Kobak KA, et al. A Novel Stable Isotope Approach Demonstrates Surprising Degree of Age-Related Decline in Skeletal Muscle Collagen Proteostasis. Funct (Oxford, England). 2021;2(4).

38. Miller BF, Reid JJ, Price JC, Lin HJL, Atherton PJ, Smith K. CORP: The use of deuterated water for the measurement of protein synthesis. J Appl Physiol. 2020;128(5):1163–1176.

39. Thomas NT, Confides AL, Fry CS, Dupont-Versteegden EE. Satellite cell depletion does not affect diaphragm adaptations to hypoxia. J Appl Physiol. 2022;133(3):637–646.

40. Wen Y, Murach KA, Vechetti IJ, et al. Myo Vision: Software for automated high-content analysis of skeletal muscle immunohistochemistry. J Appl Physiol. 2018;124(1):40–51.

41. Matsumoto T, Wakefield L, Tarlow BD, Grompe M. In Vivo Lineage Tracing of Polyploid Hepatocytes Reveals Extensive Proliferation during Liver Regeneration. Cell Stem Cell. 2020;26(1):34-47.e3.

42. Senyo SE, Steinhauser ML, Pizzimenti CL, et al. Mammalian heart renewal by pre-existing cardiomyocytes. Nature. 2013;493(7432):433–436.

43. Tanaka EM, Gann AAF, Gates PB, Brockes JP. Newt Myotubes Reenter the Cell Cycle by Phosphorylation of the Retinoblastoma Protein. J Cell Biol. 1997;136(1):155.

44. McGann CJ, Odelberg SJ, Keating MT. Mammalian myotube dedifferentiation induced by newt regeneration extract. Proc Natl Acad Sci U S A. 2001;98(24):13699–13704.

45. Pajcini K V., Corbel SY, Sage J, Pomerantz JH, Blau HM. Transient inactivation of Rb and ARF yields regenerative cells from postmitotic mammalian muscle. Cell Stem Cell. 2010;7(2):198–213.

46. Bradley LA, Young A, Li H, Billcheck HO, Wolf MJ. Loss of Endogenously Cycling Adult Cardiomyocytes Worsens Myocardial Function. Circ Res. 2021;128(2):155–168.

47. Ali SR, Hippenmeyer S, Saadat L V., Luo L, Weissman IL, Ardehali R. Existing cardiomyocytes generate cardiomyocytes at a low rate after birth in mice. Proc Natl Acad Sci U S A. 2014;111(24):8850–8855.

48. Roberts MD, Mobley CB, Vann CG, et al. Synergist ablation-induced hypertrophy occurs more rapidly in the plantaris than soleus muscle in rats due to different molecular mechanisms. Am J Physiol -Regul Integr Comp Physiol. 2020;318(2):R360–R368.

49. Øvrebø JI, Edgar BA. Polyploidy in tissue homeostasis and regeneration. Dev. 2018;145(14).

50. Xie S, Swaffer M, Skotheim JM. Eukaryotic Cell Size Control and Its Relation to Biosynthesis and Senescence. Annu Rev Cell Dev Biol. 2022;38(1).

51. Derks W, Bergmann O. Polyploidy in Cardiomyocytes: Roadblock to Heart Regeneration? Circ Res. 2020;126(4):552–565.

52. Malliaras K, Zhang Y, Seinfeld J, et al. Cardiomyocyte proliferation and progenitor cell recruitment underlie therapeutic regeneration after myocardial infarction in the adult mouse heart. EMBO Mol Med. 2013;5(2):191–209.

53. Mohamed TMA, Ang YS, Radzinsky E, et al. Regulation of Cell Cycle to Stimulate Adult Cardiomyocyte Proliferation and Cardiac Regeneration. Cell. 2018;173(1):104-116.e12.

54. Lane AN, Fan TWM. Regulation of mammalian nucleotide metabolism and biosynthesis. Nucleic Acids Res. 2015;43(4):2466–2485.

55. Fasullo M, Endres L. Nucleotide salvage deficiencies, DNA damage and neurodegeneration. Int J Mol Sci. 2015;16(5):9431–9449.

