## Supplementary data for "Skeletal muscle nuclei in mice are not post-mitotic"

Corresponding Author:

Benjamin F Miller,

**Mouse Models**: The mouse models used for breeding are provided in **Supplementary Table 1**. The satellite cell specific conditional ablation mouse (Pax7-DTA) was generated as previously described^1^ by crossing Pax7CreER/CreER and Rosa26DTA/DTA strains. To generate the adult skeletal muscle myonuclear specific GFP mouse HSArtTA;Tet-O-H2B-GFP (HSA-GFP), we crossed the muscle‐specific Tet‐On (HSA‐rtTA) mouse^2^ with the tetracycline response element histone 2b green fluorescent protein (TRE‐H2B‐GFP) mouse^3^. To generate the Pax7-DTA;HSA-GFP mouse we first crossed, through two rounds of breeding, the Pax7CreER/CreER to HSA-rtTA and the Rosa26DTA/DTA to the TetO-H2B-GFP to generate Pax7CreER/CreER; HSA-rtTA and Rosa26DTA/DTA;TetO-H2B-GFP, respectively. These two strains were then crossed to generate the Pax7-DTA;HSA-GFP mouse.

**Supplementary Table 1**: List of used animal models in the study.

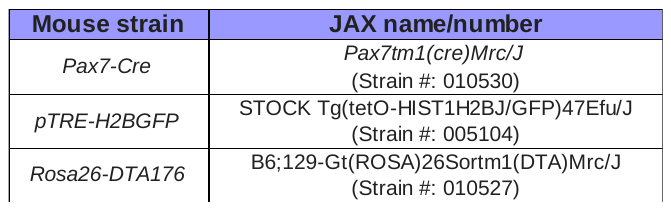

**Genotyping:** DNA was extracted from tails, and 2 μl (5-20 ng) was used in the subsequent PCR reaction. Each 25-μl PCR reaction contained Green GoTaq Master mix (Promega), 0.5 μM primers and nuclease free water. Primer sequences and used PCR conditions are provided in **Supplementary Table 2**. The Pax7-DTA;HSA-GFP: We used a two-factor validation approach, which includes SYBR based PCR followed by agarose gel PCR. Agarose gel PCR was used to confirm genotype of HSA-rtTA and H2B-GFP as described in HSA-GFP mice.

**Supplementary Table 2**: List of primer sequences and PCR conditions used for mice genotyping.

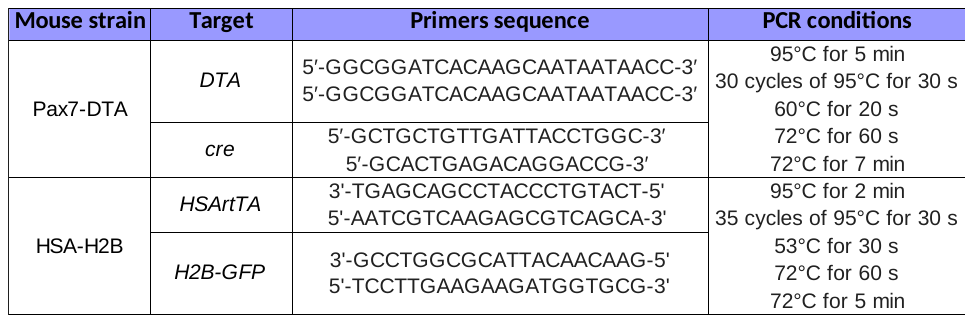

**Supplementary Table 3:** Detailed information about animals used in the study.

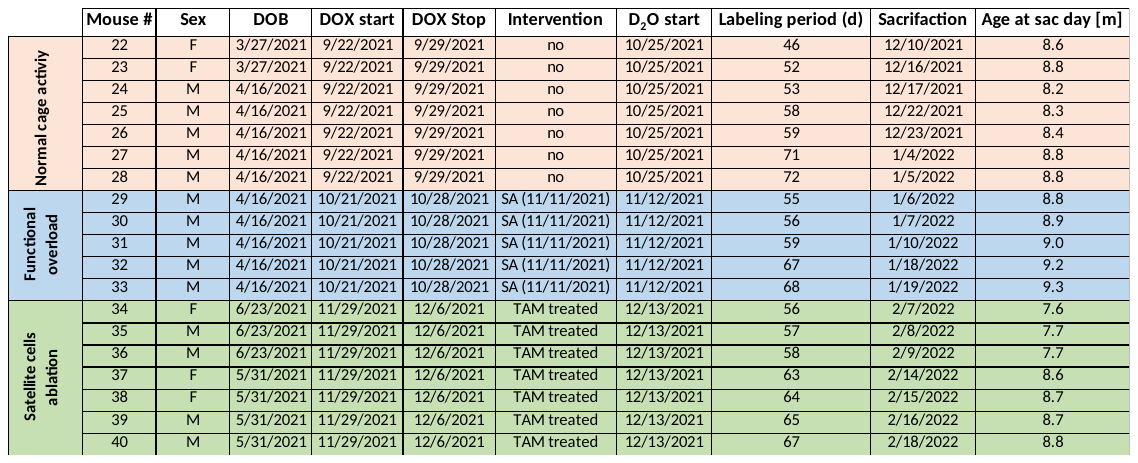

**Supplementary Table 4**: Antibodies used for immunohistochemistry analysis and validation of HSA-GFP skeletal muscles.

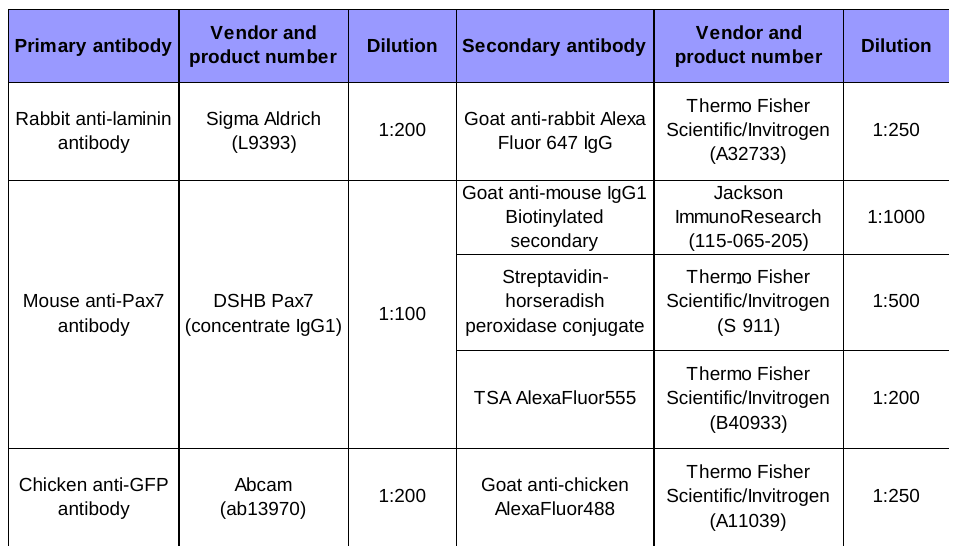

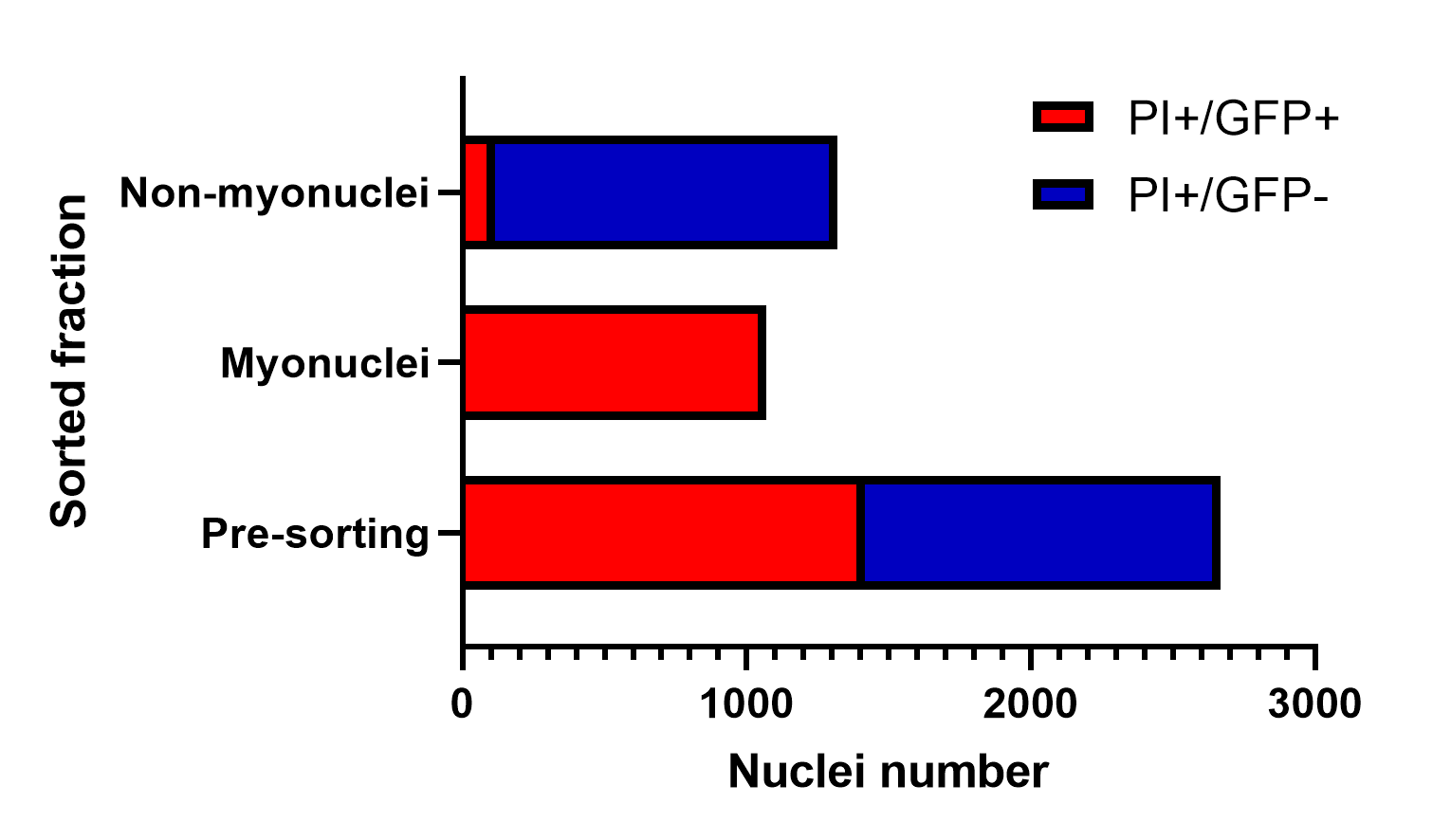

**Supplementary Figure 1: Representative quantification of the nuclei sorting efficiency.** The aliquots of crude nuclear fraction before FACS sorting, as well as sorted myonuclear and non-myonuclear fractions were analyzed for their composition. The pre-sorted fraction contained 1416 PI^+^/GFP^+^ nuclei (myonuclei) and 1252 PI^+^/GFP^-^ nuclei coming form the other cell types. After sorting, myonuclear fraction had only PI^+^/GFP^+^ nuclei (1071 counts), while non-myonuclear fraction consisted of 114 PI^+^/GFP^+^ nuclei and 1205 PI^+^/GFP^-^ nuclei.

**Supplementary Table 5: Calculations for DNA fraction synthesis rates in nuclei from HSA-H2B mice kept in normal cage conditions.**
Data were registered using unlabeled animal corrections, and bone marrow was used as a fully turned over pool.

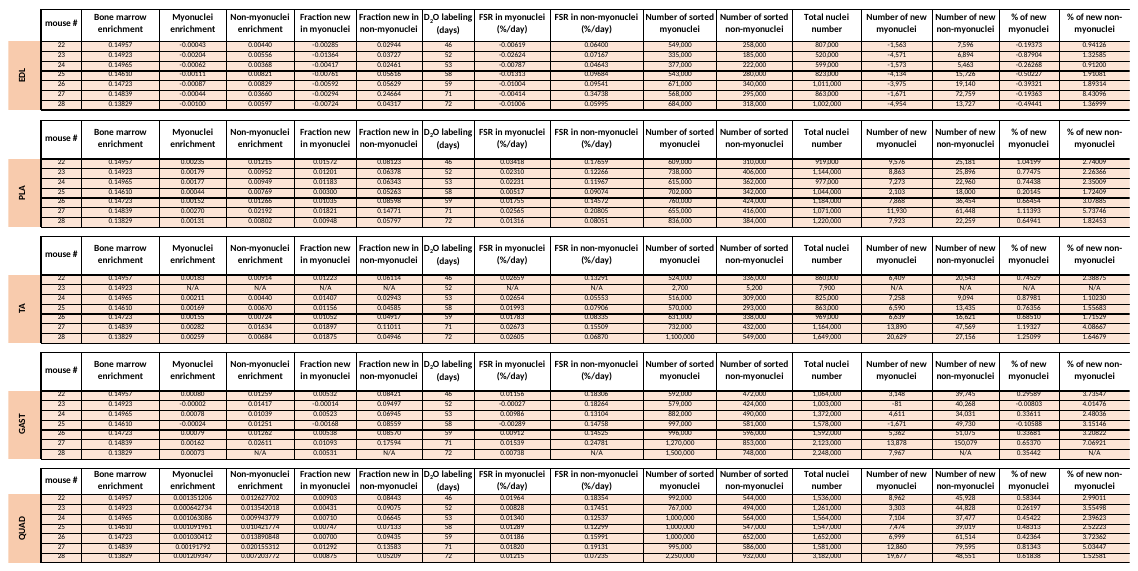

**Supplementary Table 6: Calculations for DNA fraction synthesis rates in nuclei from HSA-H2B mice kept in normal cage conditions.**
Data were registered using unlabeled animal corrections, and mass isotopomer distribution analysis (MIDA) adjustment was used.

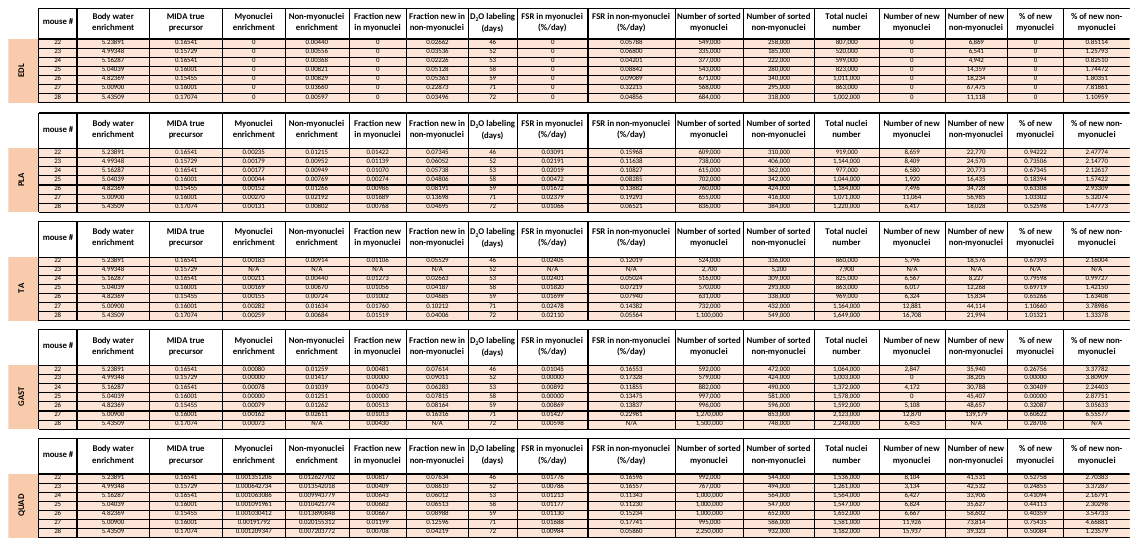

**Supplementary Table 7: Calculations for DNA fraction synthesis rates in nuclei from HSA-H2B mice after functional overload.**
Data were registered using unlabeled animal corrections, and bone marrow was used as a fully turned over pool.

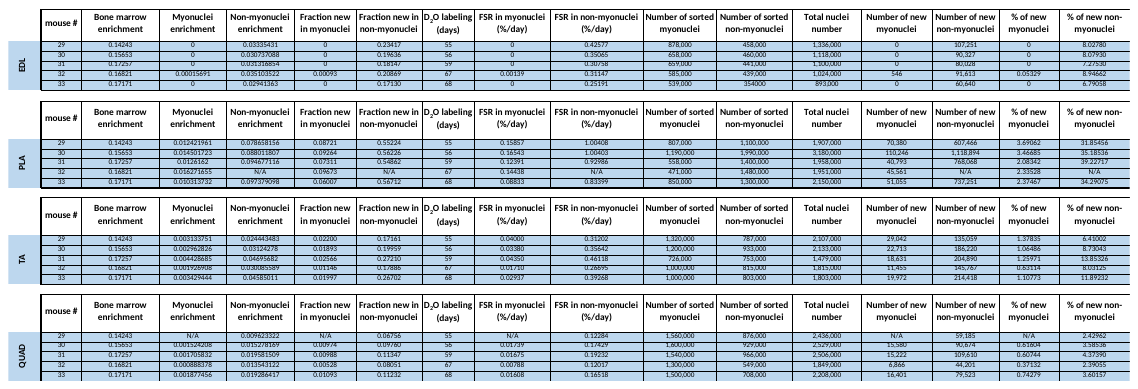

**Supplementary Table 8: Calculations for DNA fraction synthesis rates in nuclei from HSA-H2B mice after functional overload.**
Data were registered using unlabeled animal corrections, and mass isotopomer distribution analysis (MIDA) adjustment was used.

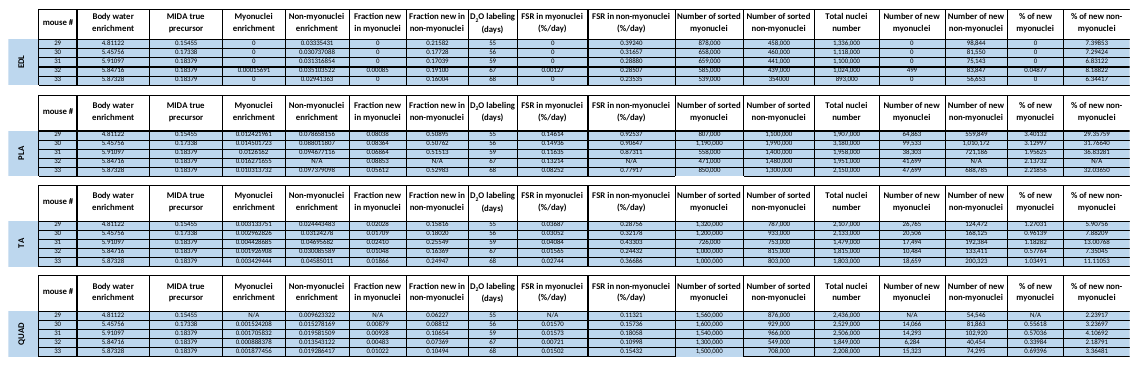

**Supplementary Table 9: Calculations for DNA fraction synthesis rates in nuclei from HSA-H2B mice after satellite cells ablation.**
Data were registered using unlabeled animal corrections, and bone marrow was used as a fully turned over pool.

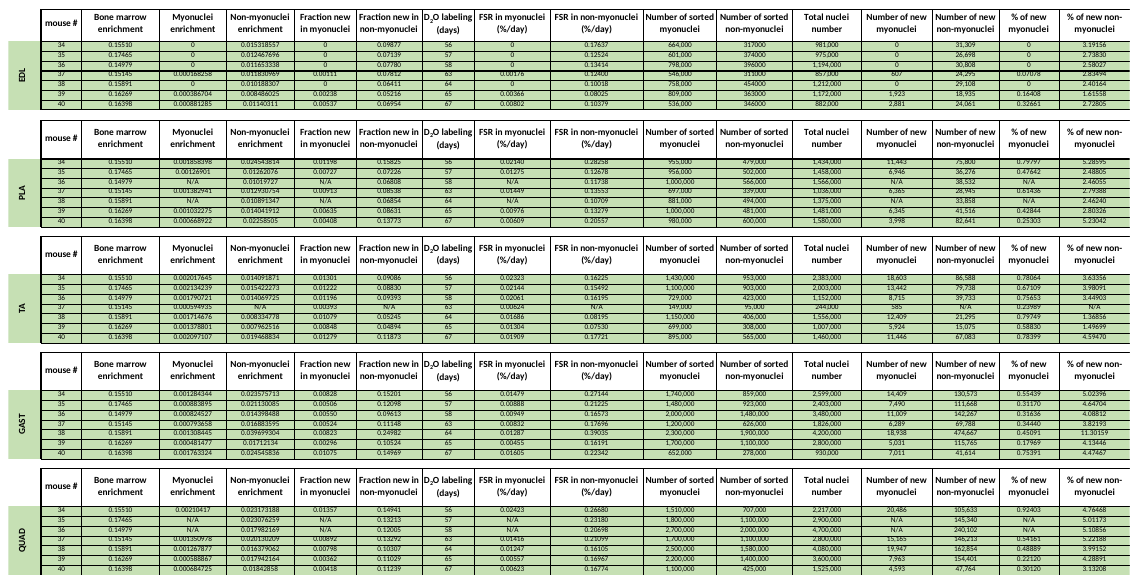

**Supplementary Table 10: Calculations for DNA fraction synthesis rates in nuclei from HSA-H2B mice after satellite cells ablation.**

Data were registered using unlabeled animal corrections, and mass isotopomer distribution analysis (MIDA) adjustment was used.

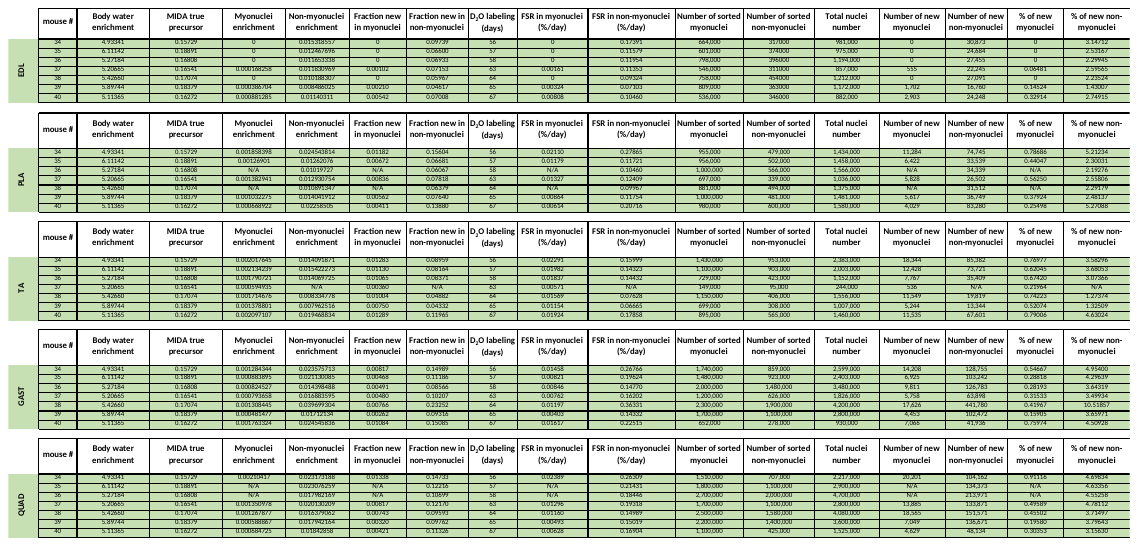

**Supplementary Table 11: Calculations for DNA fraction synthesis rates in nuclei from HSA-H2B mice kept in normal cage conditions.**
Data were registered using standard curve corrections, and bone marrow was used as a fully turned over pool.

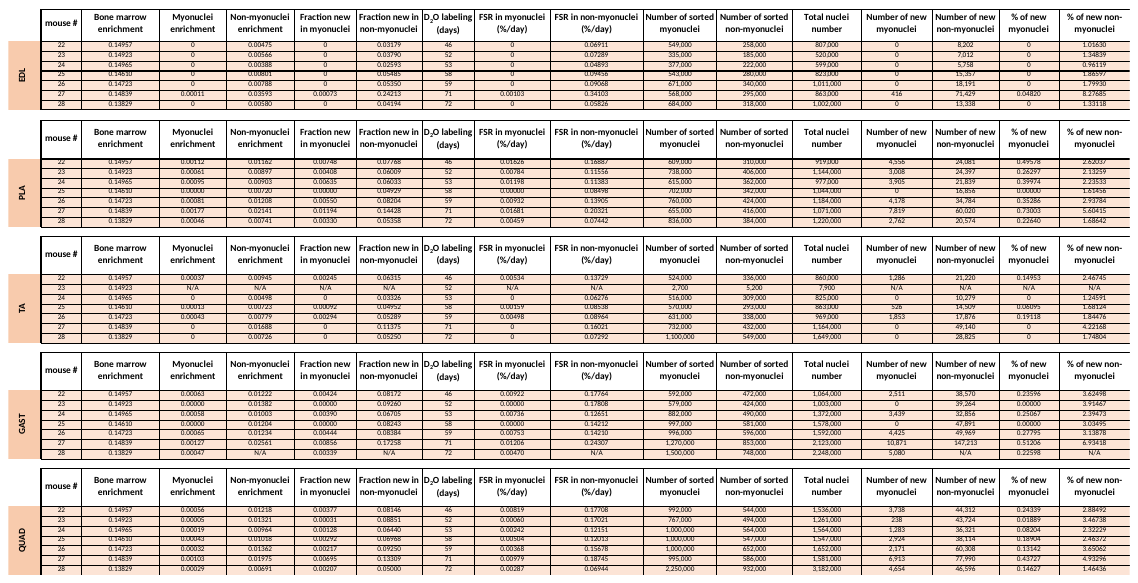

**Supplementary Table 12: Calculations for DNA fraction synthesis rates in nuclei from HSA-H2B mice kept in normal cage conditions.**
Data were registered using standard curve corrections, and mass isotopomer distribution analysis (MIDA) adjustment was used.

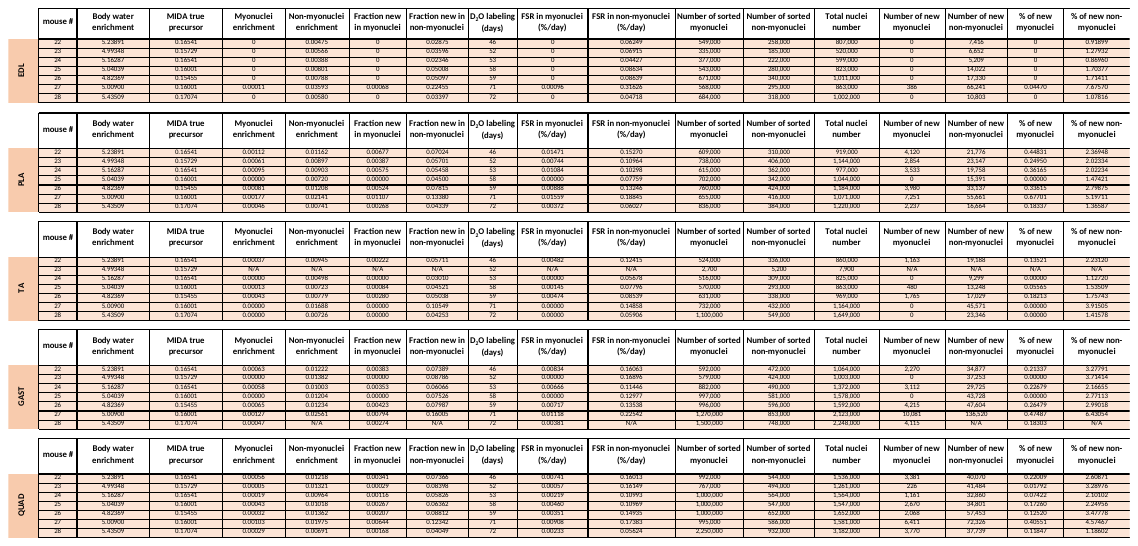

**Supplementary Table 13: Calculations for DNA fraction synthesis rates in nuclei from HSA-H2B mice after functional overload.**
Data were registered using standard curve corrections, and bone marrow was used as a fully turned over pool.

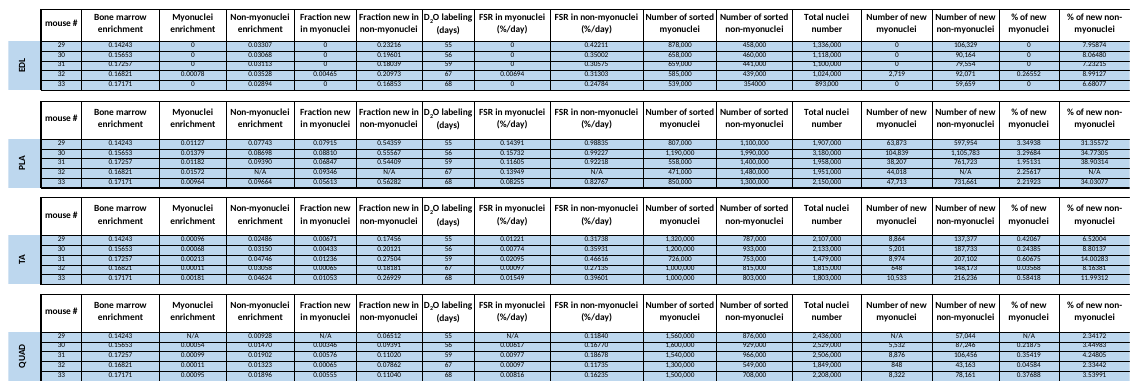

**Supplementary Table 14: Calculations for DNA fraction synthesis rates in nuclei from HSA-H2B mice after functional overload.**
Data were registered using standard curve corrections, and mass isotopomer distribution analysis (MIDA) adjustment was used.

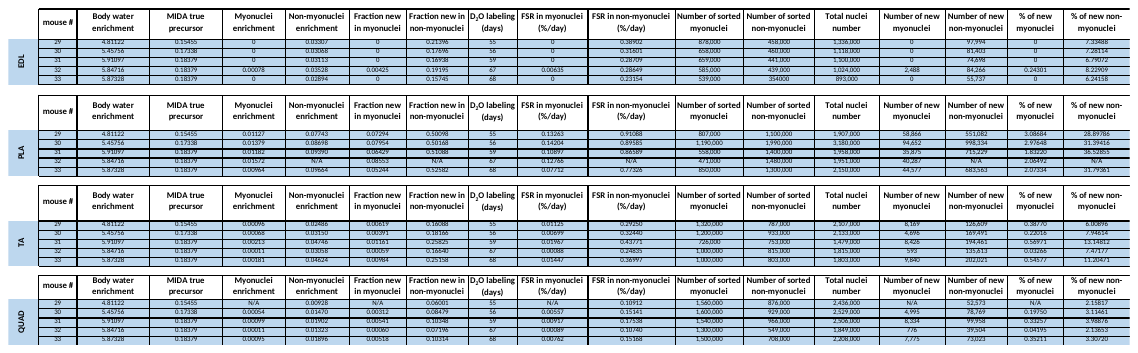

**Supplementary Table 15: Calculations for DNA fraction synthesis rates in nuclei from HSA-H2B mice after satellite cells ablation.**
Data were registered using standard curve corrections, and bone marrow was used as a fully turned over pool.

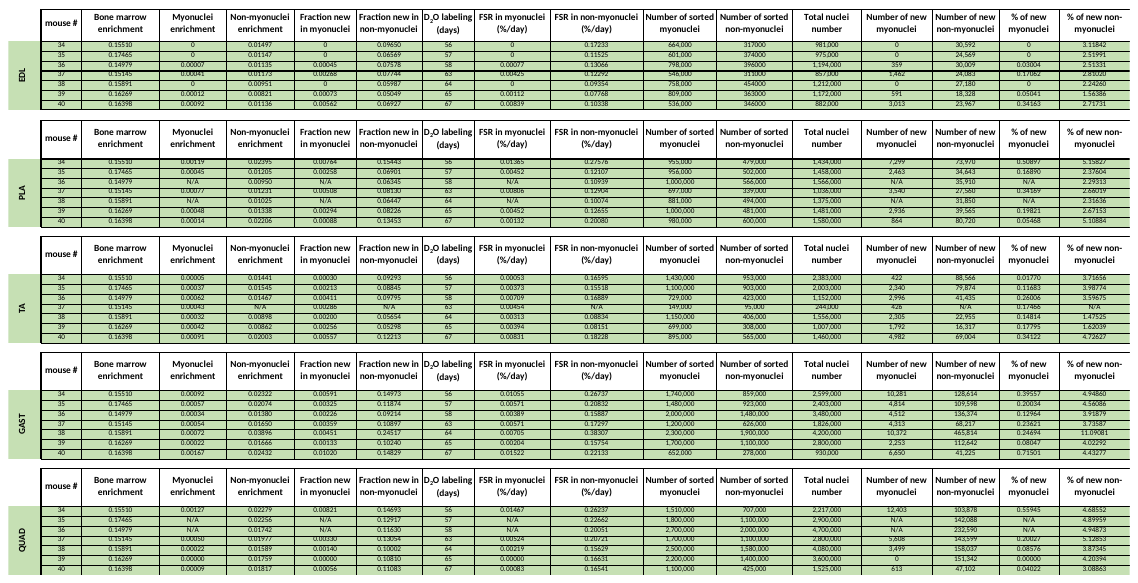

**Supplementary Table 16: Calculations for DNA fraction synthesis rates in nuclei from HSA-H2B mice after satellite cells ablation.**
Data were registered using standard curve corrections, and mass isotopomer distribution analysis (MIDA) adjustment was used.

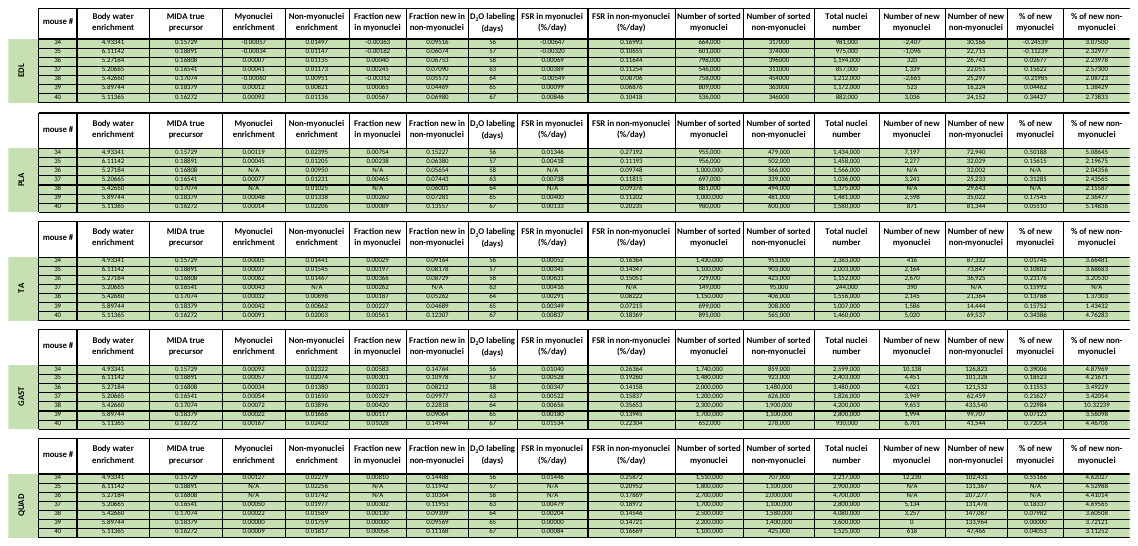

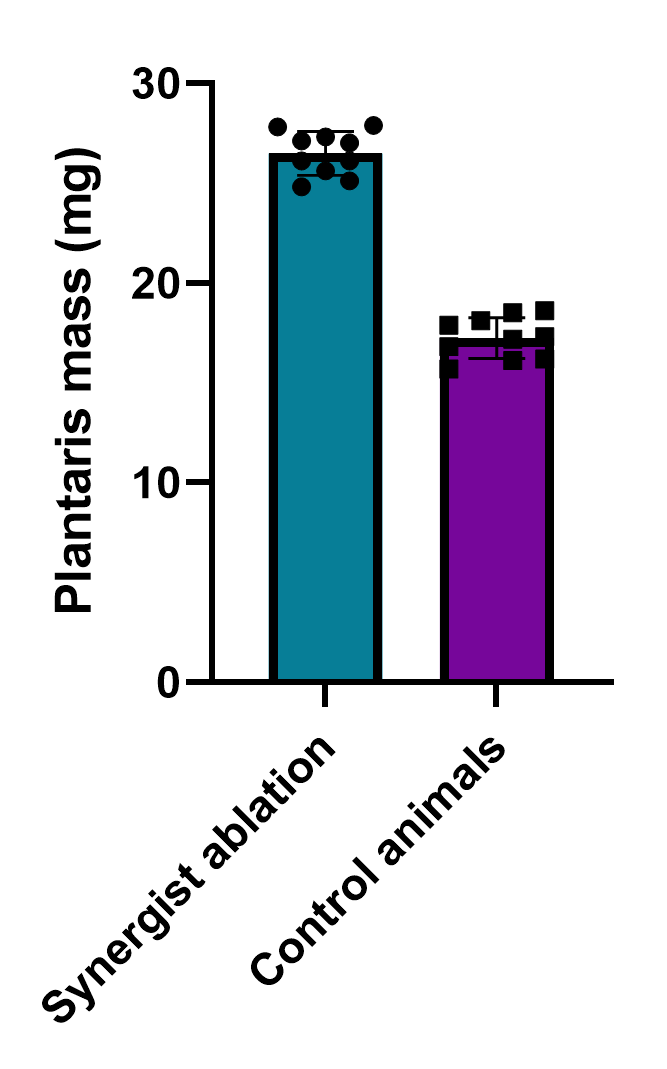

**Supplementary Figure 2: Plantaris muscle mass increased after synergist ablation when compared to control animals.** 8 weeks after synergist ablation, overloaded plantaris muscles mass from HSA-H2B mice was 26.48 ± 1.10 mg. The mass of plantaris muscles collected from age-matching control animals was 17.24 ± 1.03 mg. Values are presented as mean ± standard deviation (SD).

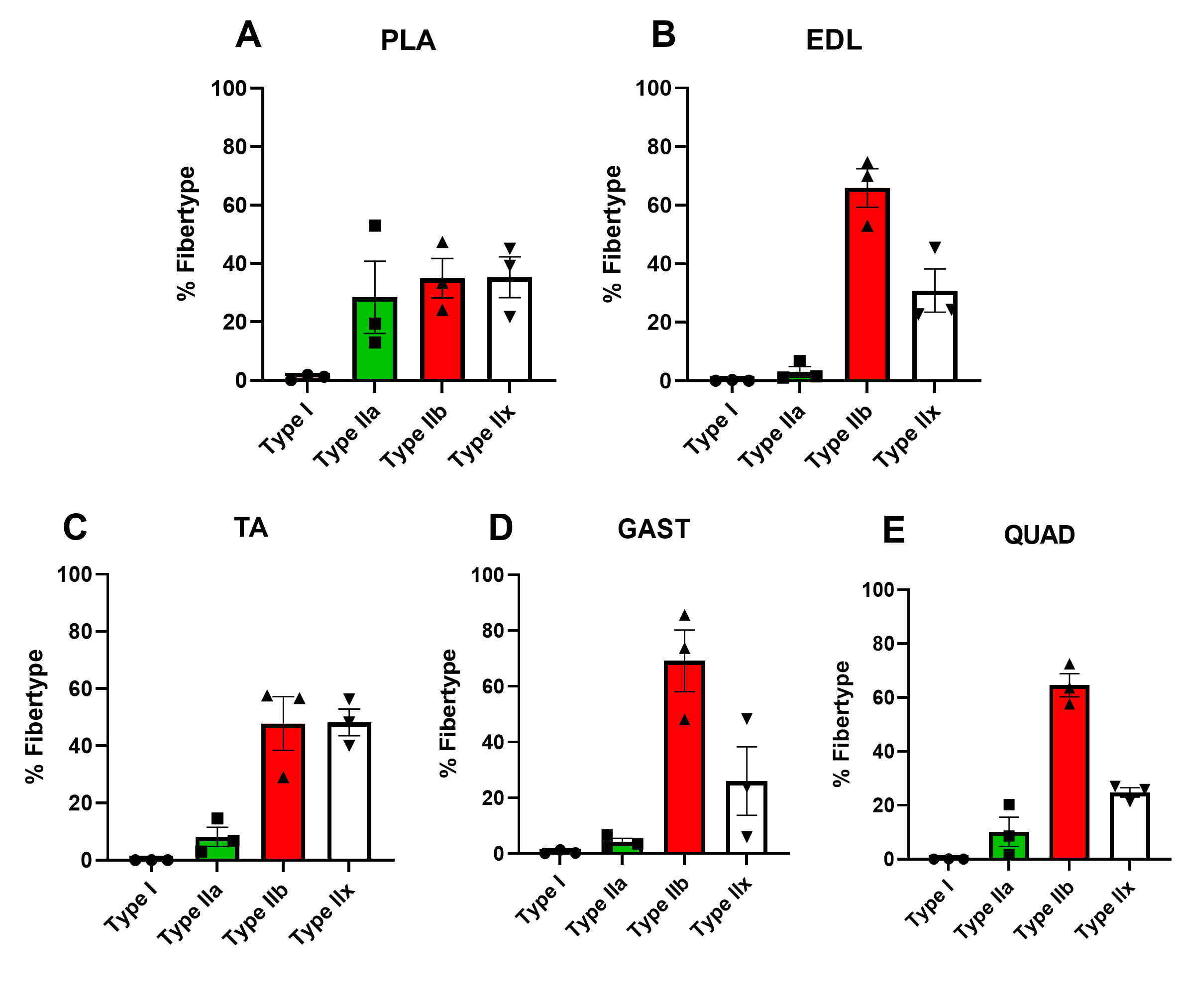

**Supplementary Figure 3: Immunohistochemical identification of HSA-H2B muscle fiber types.** Proportion of fibers in plantaris PLA **(A)**, extensor digitorum longus EDL **(B)**, tibialis anterior TA **(C)**, gastrocnemius GAST **(E)**, quadriceps QUAD **(F)** skeletal muscles.

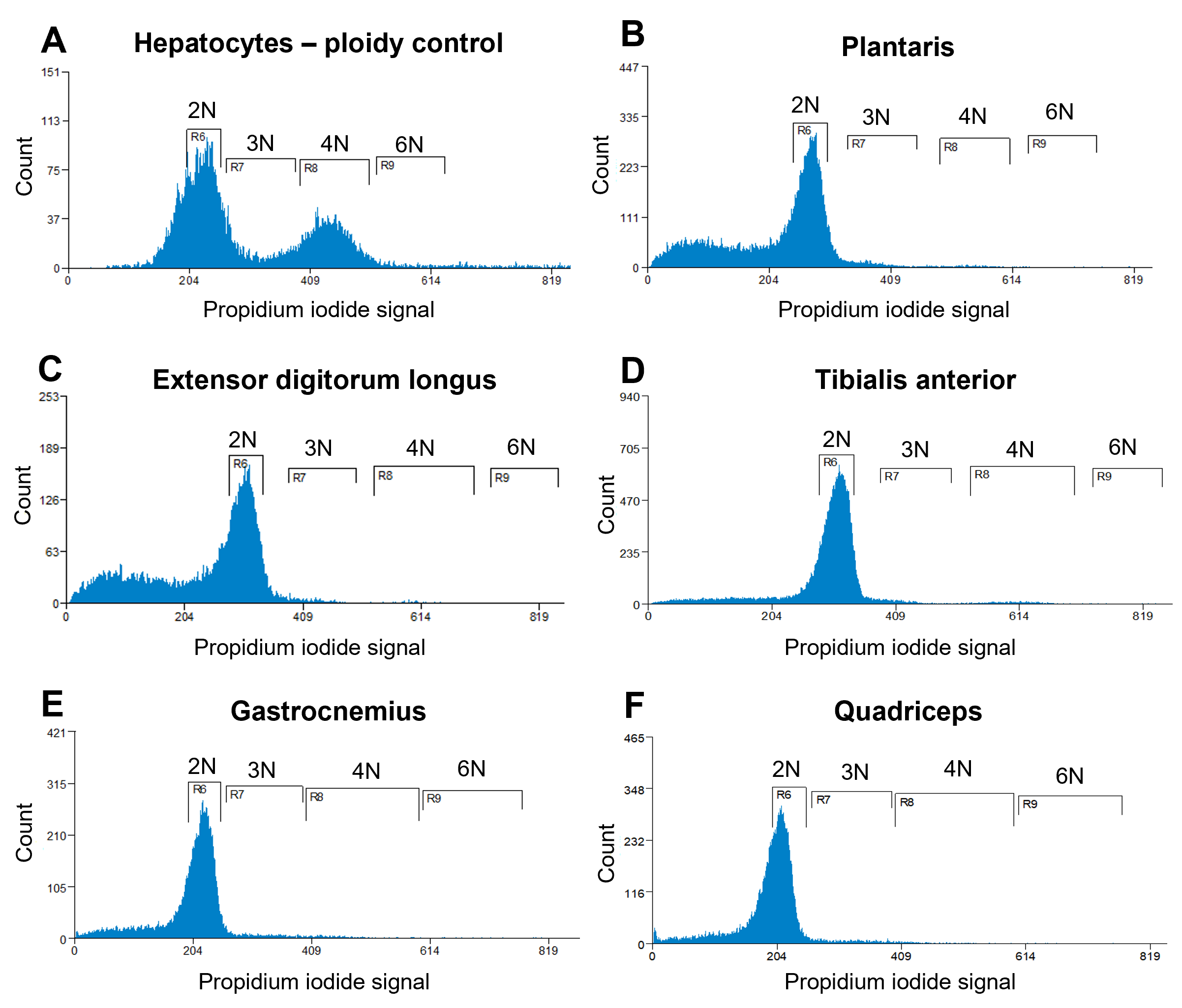

**Supplementary Figure 4: Representative analysis of ploidy levels in sorted myonuclei from muscles from animals kept in normal cage conditions.** Cell flow cytometry of hepatocytes **(A)**, myonuclei isolated from plantaris **(B)**, extensor digitorum longus **(C)**, tibialis anterior **(D)**, gastrocnemius **(E)**, quadriceps **(F)**. The peaks corresponding to diploid nuclei are labeled 2N, triploid nuclei as 3N, tetraploid nuclei as 4N and hexaploid nuclei as 6N.

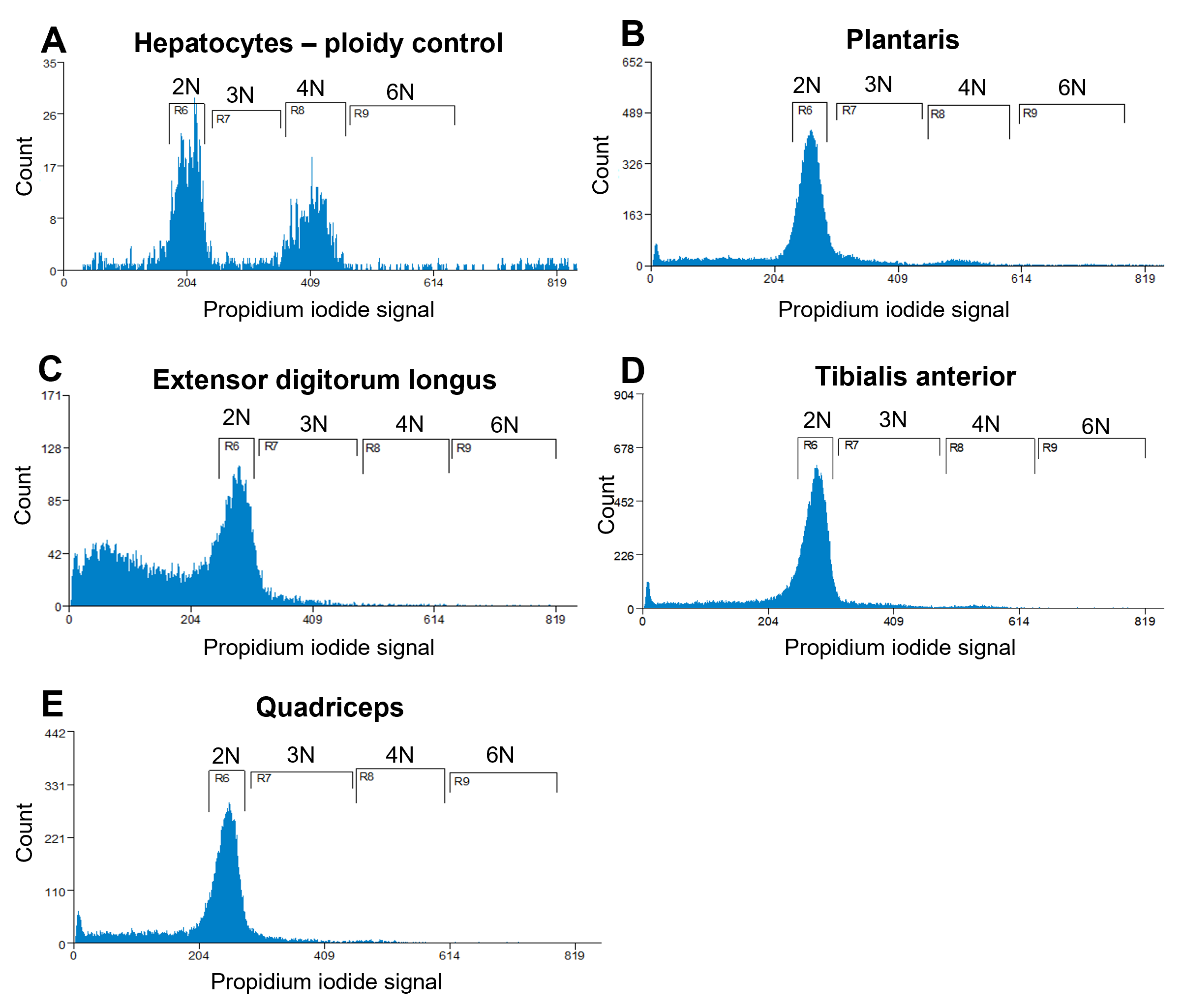

**Supplementary Figure 5: Representative analysis of ploidy levels in sorted myonuclei from muscles from animals with functional overload.** Cell flow cytometry of hepatocytes **(A)**, myonuclei isolated from plantaris **(B)**, extensor digitorum longus **(C)**, tibialis anterior **(D)**, quadriceps **(E)**. The peaks corresponding to diploid nuclei are labeled 2N, triploid nuclei as 3N, tetraploid nuclei as 4N and hexaploid nuclei as 6N.

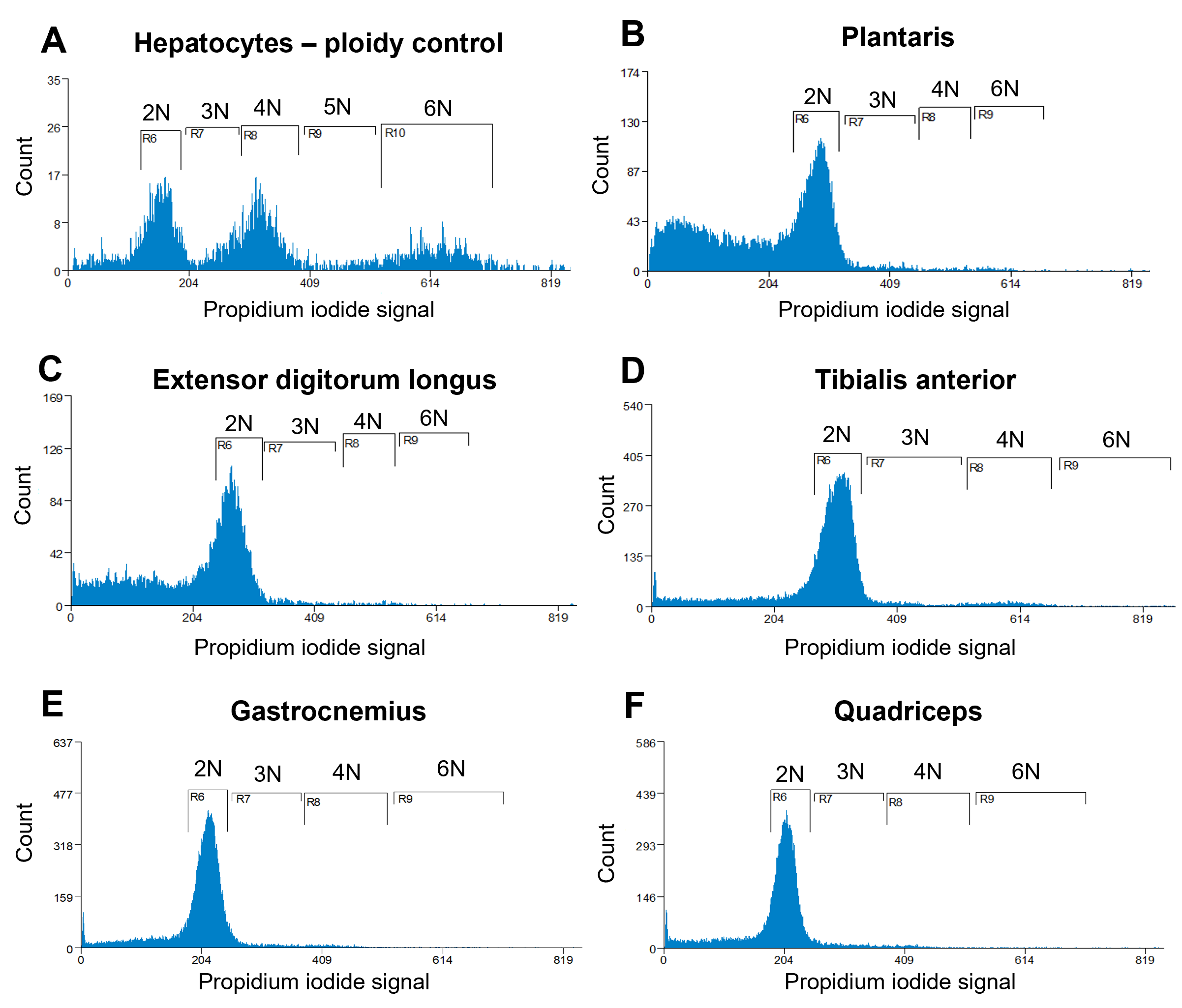

**Supplementary Figure 6: Representative analysis of ploidy levels in sorted myonuclei from muscles from animals after satellite cells ablation.** Cell flow cytometry of hepatocytes **(A)**, myonuclei isolated from plantaris **(B)**, extensor digitorum longus **(C)**, tibialis anterior **(D)**, gastrocnemius **(E)**, quadriceps **(F)**. The peaks corresponding to diploid nuclei are labeled 2N, triploid nuclei as 3N, tetraploid nuclei as 4N, pentaploid nuclei as 5N and hexaploid nuclei as 6N.

**References**

1.	Mccarthy JJ, Mula J, Miyazaki M, et al. Effective fiber hypertrophy in satellite cell-depleted skeletal muscle. *Development*. 2011;138(17):3657-3666.

2.	Iwata M, Englund DA, Wen Y, et al. A novel tetracycline-responsive transgenic mouse strain for skeletal muscle-specific gene expression. *Skelet Muscle*. 2018;8(1):1-8.

3.	Tumbar T, Guasch G, Greco V, et al. Defining the epithelial stem cell niche in skin. *Science*. 2004;303(5656):359-363.
